# Tandem sulfofucolytic-sulfolactate sulfolyase pathway for catabolism of the rare sulfosugar sulfofucose

**DOI:** 10.1101/2023.08.01.551581

**Authors:** Adam W. E. Stewart, Jinling Li, Mihwa Lee, Jessica M. Lewis, Marion Herisse, Vinzenz Hofferek, Malcolm J. McConville, Sacha J. Pidot, Nichollas E. Scott, Spencer J. Williams

**Affiliations:** School of Chemistry and Bio21 Molecular Science and Biotechnology Institute, University of Melbourne, Parkville, Victoria 3010, Australia; Department of Microbiology and Immunology, University of Melbourne, at the Peter Doherty Institute for Infection and Immunity, Victoria 3000, Australia; Department of Biochemistry and Pharmacology, Bio21 Molecular Science and Biotechnology Institute, The University of Melbourne, Parkville, Victoria, Australia

**Author notes:** Address correspondence to Spencer J. Williams. Equal first authors (Adam W. E. Stewart, Jinling Li).

**Keywords:** organosulfur, metabolism, sulfoglycolysis, sulfur cycle, catabolism. Running title: Sulfofucose catabolism pathway in *Paracoccus wurundjeri*

## Abstract

The rare sulfosugar sulfofucose (6-deoxy-6-sulfo-D-galactose) is catabolized by *Paracoccus wurundjeri sp. nov.* strain Merri via a novel tandem pathway that couples sulfofucolytic and sulfolactate sulfolyase reactions. We isolated *P. wurundjeri* from soil and demonstrated its ability to utilize SFuc as the sole carbon source, leading to complete degradation of sulfofucose and release of sulfite. Proteomics and metabolomics analysis revealed a sulfofucolytic variant of the Entner–Doudoroff (ED) pathway, wherein sulfofucose is converted to sulfolactaldehyde through oxidation, lactone hydrolysis, dehydration and retro-aldol steps. The sulfolactaldehyde is subsequently degraded to pyruvate and sulfite via a downstream biomineralization pathway involving sulfolactate as an intermediate. *In vitro* reconstitution of the key enzymatic steps validated the pathway, identifying four enzymes: SfcH (dehydrogenase), SfcD (lactonase), SfcF (dehydratase), and SfcE (aldolase), that collectively convert sulfofucose to sulfolactaldehyde and pyruvate. Our findings provide insight into microbial utilization of rare sulfosugars and expand the known metabolic versatility of the genus *Paracoccus*.

**Importance:** Sulfosugars, such as sulfoquinovose, are integral to the global sulfur cycle, but the metabolism of rarer sulfosugars, like sulfofucose, remains poorly understood. This study identifies a unique bacterial catabolic pathway for sulfofucose degradation, comprising a sulfofucolytic ED pathway followed by biomineralization to sulfite. The identification of this tandem pathway highlights the metabolic flexibility of bacteria to adapt carbohydrate catabolic pathways for processing structurally similar sulfosugars. This work sheds light on the ecological role of sulfofucose metabolism and broadens our understanding of bacterial sulfur metabolism, offering potential implications for biogeochemical sulfur cycling and biotechnological applications.

## Introduction

Sulfur is an essential macronutrient and is incorporated into a wide range of organosulfur compounds that serve as metabolic currencies and facilitate sulfur exchange between producer and consumer organisms (1, 2). The biosynthesis and catabolism of these organosulfur metabolites form intricate biochemical networks within the global sulfur cycle (3, 4). Organosulfur compounds of global significance include the sulfur-containing amino acids cysteine and methionine, the osmoprotectants dimethylsulfoniopropionate (5) and dimethyloxosulfonium propionate (6), the short-chain organosulfonates 2,3-dihydroxypropanesulfonate (DHPS) and sulfolactate (SL) (7, 8), and plant and cyanobacterial sulfolipid (sulfoquinovosyl diglycerol; SQDG), which contains the sulfosugar sulfoquinovose (SQ, 6-deoxy-6-sulfo-D-glucose) (9).

Over the past three decades, extensive work has elucidated the biosynthetic route to SQ in plants and cyanobacteria, as well as the diverse bacterial pathways used for its degradation (**Figure 1a**) (9–12). Glycoside hydrolases known as sulfoquinovosidases initiate catabolism by cleaving SQ from glycoconjugates such as SQDG and sulfoquinovosyl glycerol (13–15). SQ breakdown then occurs via multiple sulfoglycolytic and sulfolytic pathways. In sulfoglycolytic bacteria, SQ is partially degraded into short-chain C2 and C3-organosulfonates (16–20), which are excreted and subsequently utilized by secondary consumer organisms (17, 21–23). These secondary organisms—facultative anaerobes or aerobes—cleave the C–S bond using sulfolyases and release sulfite to enable carbon assimilation (24, 25). In anaerobic environments, short-chain organosulfonates serve as terminal electron acceptors or sulfur sources, a process facilitated by glycyl radical enzymes (26). In contrast, sulfolytic bacteria can directly cleave the C–S bond in SQ, releasing sulfite and converting the entire carbon backbone to glucose for entry into central carbon metabolism (19, 27).

**Figure 1.**
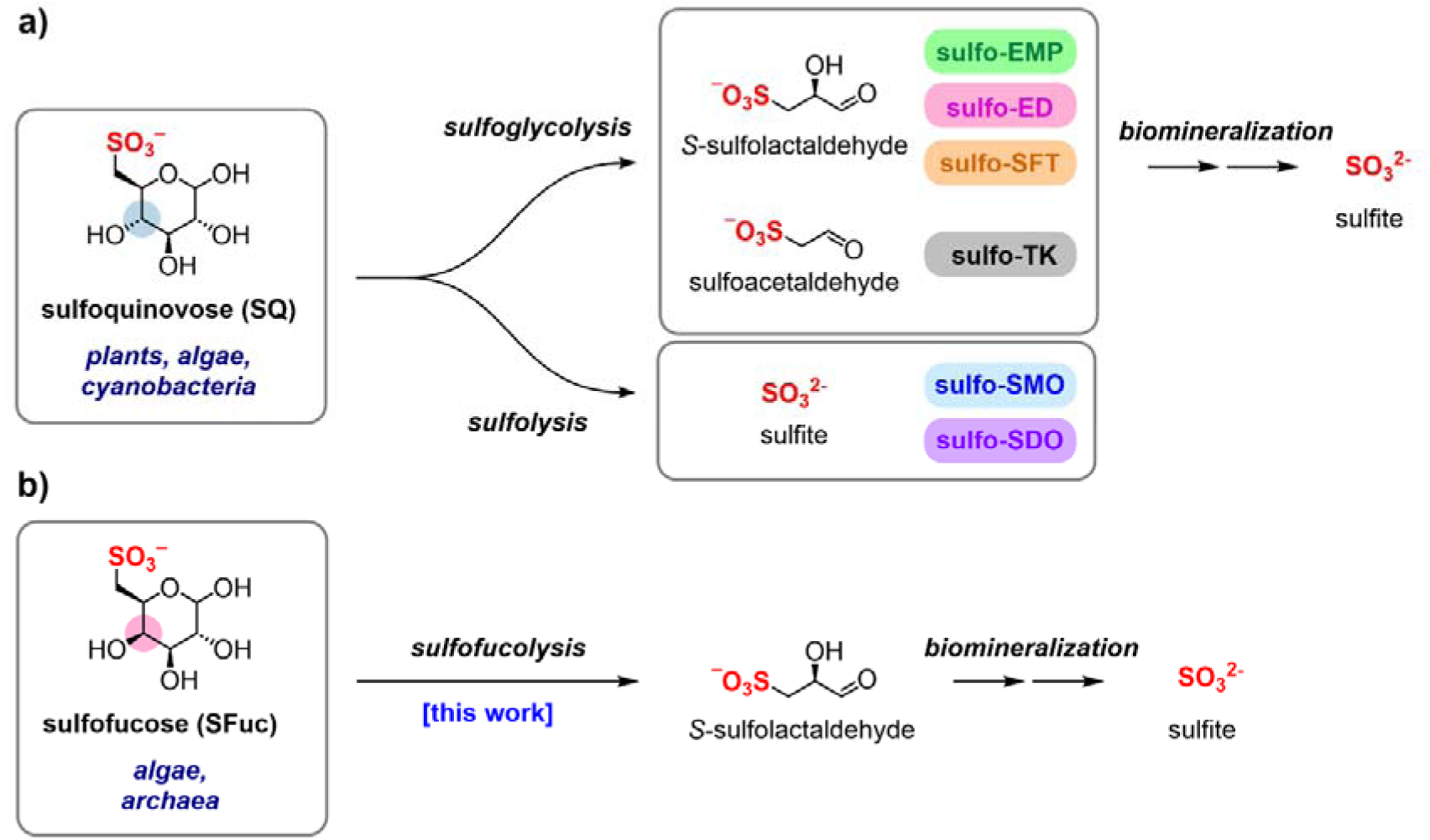
Breakdown of the sulfosugars sulfoquinovose and sulfofucose. (a) Degradation of sulfoquinovose to sulfoacetaldehyde (SA) or sulfolactaldehyde (SLA) by sulfoglycolysis (sulfoglycolytic Embden−Meyerhof−Parnas (sulfoEMP); sulfoglycolytic Entner−Doudoroff (sulfo-ED); sulfoglycolytic sulfofructose transaldolase (sulfo-SFT); sulfoglycolytic transketolase (sulfo-TK) pathways) or sulfolysis to sulfite (sulfolytic SQ monooxygenase (sulfo-SMO); and sulfolytic SQ dioxygenase (sulfo-SDO) pathways). (b) Sulfofucose has been identified in glycoconjugates from algae and archaea, this work discloses a two-phase sulfofucolysis−biomineralization breakdown pathway.

While SQ metabolism is well-characterized, much less is known about other sulfosugars. One such compound is sulfofucose (SFuc, 6-deoxy-6-sulfo-D-galactose), a rare sulfosugar reported in the glycolipids of algae from the order *Fucacaea*, which are distributed across the North Atlantic Ocean (28), and in cell surface N-linked glycans from the extremophile archaeon *Thermoplasma acidophilum* (**Figure 1b**) (29). SFuc differs from sulfoquinovose only in the configuration of carbon-4. Despite this minimal structural difference, no biosynthetic or catabolic pathways for SFuc have been identified.

Here, we report the isolation of *Paracoccus wurundjeri sp. nov.* strain Merri, a bacterium capable of using SFuc as its sole carbon source. Using uniformly ^13^C-labeled SFuc, we show that the entire carbon skeleton is metabolized and that sulfite is released as the end-product. Comparative proteomics reveal increased abundance of proteins associated with organosulfur metabolism during growth on SFuc. Metabolomics analysis of cell lysates identified a pathway intermediate consistent with an Entner–Doudoroff (ED) like pathway for SFuc metabolism to sulfolactaldehyde (SLA), and a sulfolyase pathway for biomineralization via SL. Finally, recombinant expression and *in vitro* reconstitution of four enzymes allowed demonstration of the individual steps of a sulfofucolytic ED pathway that converts SFuc to SLA and pyruvate. These findings support the operation of a tandem pathway comprising oxidative cleavage of SFuc to SLA, followed by canonical biomineralization to sulfite via SL sulfolyase.

## Results

### Isolation and characterization of a *Paracoccus wurundjeri* bacterium able to grow on sulfofucose

A soil sample from Joe’s Garden, an inner-Melbourne market garden that has been continuously farmed by Chinese and Italian gardeners for 150 years, was used to inoculate a vitamin-enriched defined media containing 10 mM SFuc as the sole carbon source. Sequential subculturing, along with repeated plating onto SFuc-containing media agar plates, followed by regrowth in SFuc liquid media led to isolation of strain Merri. This strain grew on glucose and SFuc to similar peak absorbance, but did not grow on SQ (**Figure 2a**). Genomic DNA was extracted from strain Merri and was sequenced using the Illumina NextSeq platform. Analysis of the 16S rRNA gene against the NCBI 16S rRNA database revealed the most similar sequence to be from *Paracoccus methylarcula* strain H1 (97.18% identity). With 97% 16S rRNA sequence identity generally recognized as the cutoff for an organism to be considered the same species, this places strain Merri firmly within the genus *Paracoccus,* but close to being considered a new species. In light of this, we performed further in silico analysis using DNA–DNA hybridization against the Genome Taxonomy Database. This analysis showed strain Merri had 94.37% average nucleotide identity (ANI) to the most closely related reference genome (*Paracoccus* sp. 003591515), which is below the threshold to be considered the same species (95% ANI) (30, 31). Given ANI is generally considered a more robust method to define taxonomic lineages (30, 32) we propose that strain Merri represents a new member of the genus *Paracoccus*, which we have designated *Paracoccus wurundjeri sp. nov.* (Strain ID: Merri), based on its isolation from the banks of Merri creek, which runs through the lands of the indigenous Wurundjeri people. General features of the draft genome are shown in **Table 1**.

**Figure 2.**
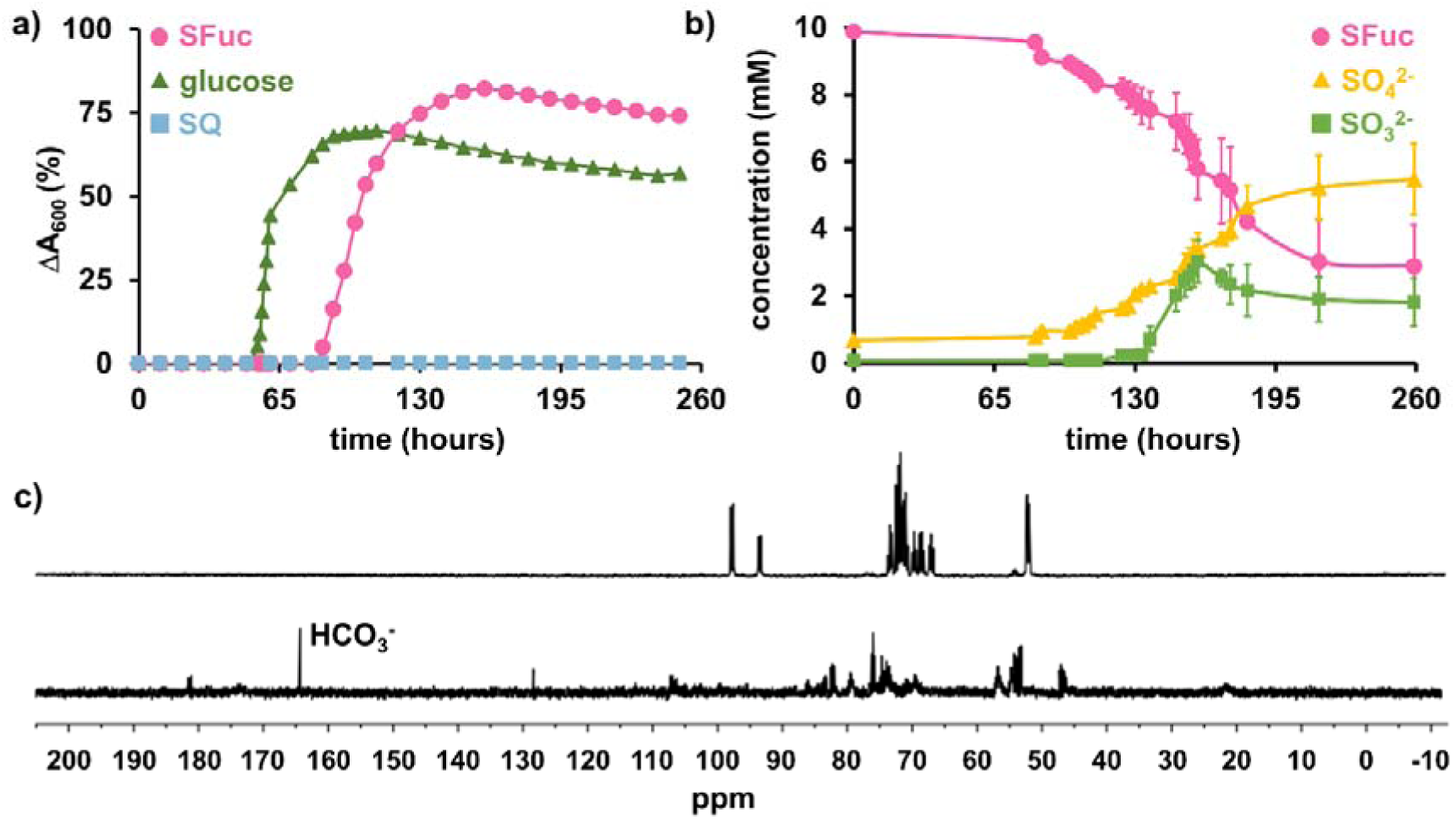
*P. wurundjeri* strain Merri grows on SFuc as a sole carbon source. (a) Growth of strain Merri on glucose, SFuc, and SQ. Turbidity was measured at A_600_ nm, and cells were cultured in M9 minimal media containing 5 mM of each carbon source, supplemented with a vitamin mix and trace metals. (b) Strain Merri uses SFuc as a carbon source and produces sulfite. Independent replicates show the change in concentration of SFuc (open circles), sulfite (open squares) and of sulfate (closed triangles) throughout growth of strain Merri on SFuc. Analyses were performed in duplicate, and errors are reported as standard deviations. (c) ^13^C NMR spectra of M9 minimal media containing 10 mM ^13^C_6_-SFuc (top) and after 5 days of growth of strain Merri at the mid-log phase (bottom).

**Table 1.** Key features of *Paracoccus wurundjeri* strain Merri (GenBank accession: Bioproject PRJNA993841) draft genome assembly.

|  | **Strain Merri** |
| --- | --- |
| Genome size | 5,082,950 |
| Number of contigs | 74 |
| GC % | 60.1 |
| Number of ORFs | 4913 |
| Number of putative genes | 4966 |
| Number of putative tRNA | 49 |
| Number of putative rRNA | 3 |
| Number of putative tmRNA | 1 |
| Number of genes assigned a function (%) | 2847 (57%) |

To trace SFuc catabolism, we examined the fate of its carbon and sulfur components upon metabolism by *P. wurundjeri* strain Merri. To examine the fate of sulfur, spent culture media from strain Merri grown on SFuc was analyzed. SFuc consumption was tracked using a reducing sugar assay, while sulfite and sulfate production were quantified with colorimetric and turbidometric assays, respectively. Most SFuc was consumed over 200 h, accompanied by the production of sulfate and sulfite (**Figure 2b**). Sulfite levels peaked at 160 hours before declining, a pattern consistent with the known oxygen sensitivity of sulfite (27). This suggests that sulfite is initially produced and subsequently undergoes autooxidation to sulfate. ^13^C NMR analysis of the culture medium of *P. wurundjeri* strain Merri grown to mid log-phase (5 d) on ^13^C_6_-SFuc revealed complete consumption of the substrate, and production of ^13^C-bicarbonate (**Figure 2c**); extended growth to stationary phase (14 d) revealed no distinct carbon-containing compounds, indicating complete utilization of the complete carbon chain (data not shown).

### Identification of genetic loci expressed upon growth on sulfofucose

To identify proteins potentially involved in SFuc metabolism, we cultured *P. wurundjeri* strain Merri using either SFuc or glucose as the sole carbon source and analyzed the resulting changes in the proteome (**Figures 3a, b**). Growth on SFuc led to significant increases in abundance of proteins primarily encoded within three genomic regions, designated clusters 1–3.

**Figure 3.**
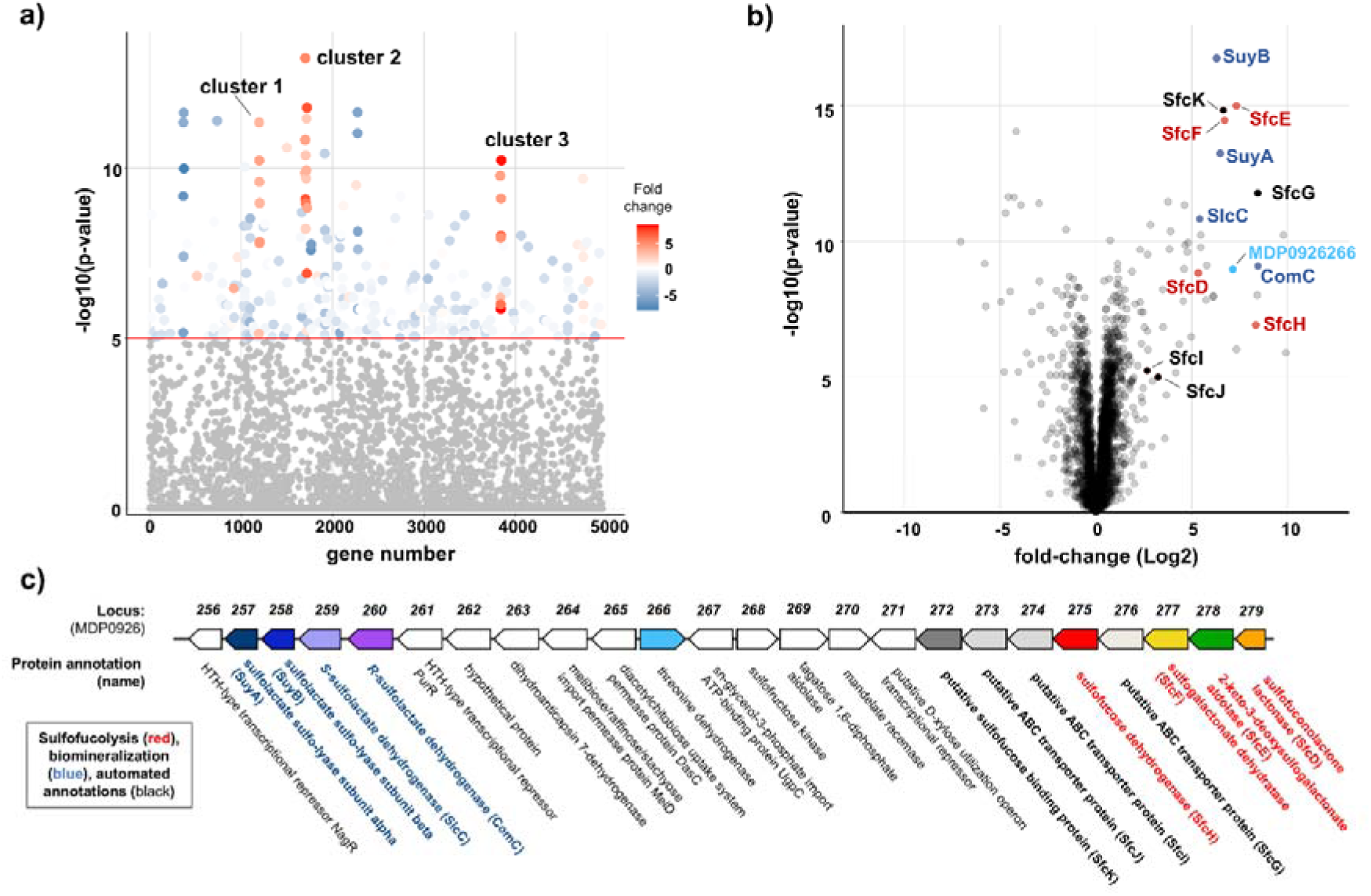
Genome and proteome analysis reveals a gene cluster for organosulfonate metabolism in *P. wurundjeri* strain Merri. Cells were cultured in vitamin-supplemented minimal media with either SFuc or glucose, and analyzed via comparative proteomics. (a) Manhattan plot highlighting three differentially expressed gene clusters. (b) Volcano plot showing proteins identified in the SFuc degradation pathway (red, SfcD, SfcE, SfcF, SfcG; blue, SlcC, ComC, SuyAB; black, putative SFuc binding protein and ABC transporter cassette). (c) Gene neighborhood diagram of "cluster 2", associated with organosulfonate metabolism. Genes annotated in red are sulfofucolytic genes; in blue are putative biomineralization genes; in bold black text are a putative SFuc binding protein and ABC transporter cassette; and in plain black text are automated genome annotations.

Cluster 1 (Locus IDs MDP0925965–0925971) comprises seven genes, all of which showed increased protein abundance (**Figure S1**). These genes encode enzymes annotated for roles in gluconeogenesis and the pentose phosphate pathway. Sulfoglycolysis circumvents the glycolytic steps that generate glucose-6-phosphate and fructose-6-phosphate—key intermediates required for the pentose phosphate pathway and the biosynthesis of sugar nucleotides and cell wall components. Previous studies in *E. coli* have shown that sulfoglycolysis via the sulfo-EMP pathway elevates gluconeogenic flux due to the generation of the C3-phosphate DHAP (33); similarly, a sulfofucolytic pathway in *P. wurundjeri* would require gluconeogenesis to supply hexose-phosphate intermediates from DHAP or pyruvate. Cluster 3 (MDP0926751–0926761) includes nine proteins with significantly increased abundance, associated with fumarate and tartrate metabolism, which are likely linked to the Krebs cycle (**Figure S1**). This cluster also harbors gene MDP0926757, annotated as a hypothetical protein but homologous to the TauE transmembrane sulfite exporter (34).

Cluster 2 (MDP0926256–0926279) contains 24 genes, many of which were increased in abundance in SFuc-grown cells (**Figure 3c, Table S1**). Some of these genes are annotated with functions related to organosulfonate metabolism, and belong to protein families (PFAM) associated with SL biomineralization in *Chromohalobacter salexigens* (25), *Paracoccus pantotrophus* (35) and *Desulfvibrio* sp. DF1 (36) (**Table S2**). Among the proteins of increased abundance were homologues of SL dehydrogenases SlcC (MDP0926259) and ComC (MDP0926260), which could act as a two-component SL racemase (**Figures S5, S6**) (25). Additionally, MDP0926258, annotated as SL sulfolyase B (SuyB), and MDP0926257, encoding a short protein homologous to SL sulfolyase A (SuyA) (**Figures S7, S8**) (35, 36), were increased in abundance. Finally, protein MDP0926266, annotated as threonine dehydrogenase, was present in increased abundance, and is a candidate for an SLA dehydrogenase for oxidation of SLA to SL. Together, these proteins may form a C3-organosulfonate biomineralization pathway that involves oxidation of *S*-sulfolactaldehyde to *S*-sulfolactate, catalyzed by MDP0926266, racemization to *R*-sulfolactate catalyzed by SlcC/ComC, and SuyAB catalyzed cleavage of *R*-sulfolactate to yield pyruvate and sulfite, with the latter exported via TauE.

Turning to the initial steps of SFuc metabolism, several genes within cluster 2 are homologous to those involved in known sulfosugar breakdown pathways. Two groups of interest were identified. The first group includes homologues of sulfo-EMP pathway proteins: MDP0926268 and MDP0926269, belonging to PF00294 and PF01791, respectively, protein families encompassing sulfofructose kinase and sulfofructose-1-phosphate aldolase (**Table S3**) (17). Possibly, these could act on the 4-epimeric SFuc-derived intermediates, such as the sulfoketohexose sulfotagatose (6-deoxy-6-sulfo-D-tagatose) and sulfotagatose-1-phosphate, with the latter enzyme MDP0926269 potentially producing SLA and DHAP. While no aldose–ketose isomerase is encoded within this cluster, a homologue is encoded elsewhere in the genome (protein MDP0925592) belonging to protein family PF07221, the same family as SQ-sulfofructose isomerase within the sulfo-EMP pathway (17). However, none of these three proteins showed significantly increased abundance in SFuc-grown cells.

The second group of genes of interest from cluster 2 includes MDP0926275, MDP0926278, and MDP0926279—the corresponding proteins of which were all increased in SFuc-grown cells and are associated with PF13561, PF03328, and PF09450, respectively. These protein families include enzymes from the sulfoglycolytic ED pathway: SQ dehydrogenase, 2-keto-3-deoxysulfogluconate (KDSG) aldolase, and sulfogluconolactone (SGL) lactonase, respectively (**Table S4**) (18). A fourth enzyme required for a complete sulfoglycolytic ED pathway, SG dehydratase (belonging to PF00920), was not encoded within cluster 2 or elsewhere in the genome. However, the protein encoded by MDP0926279 in cluster 2 was increased in abundance and belongs to protein families PF13378 and PF02746, families to which D-galactonate dehydratase belongs. Notably, D-galactonate bears close structural resemblance to sulfogalactonate (SGal). Thus, we propose that protein MDP0926279 may be capable of dehydrating a sulfofucolytic ED intermediate such as SGal. Associated with these genes was a likely ABC transporter cassette and substrate binding protein, which we assign as a putative SFuc importation system.

Collectively, these data support the existence of either an EMP-like or ED-like sulfofucolysis pathway. In the proposed sulfofucolytic EMP pathway, SFuc would be isomerized to sulfotagatose (by protein MDP0925592), phosphorylated to sulfotagatose-1-phosphate (by protein MDP0926268), and then cleaved via a retroaldol reaction (by protein MDP0926269) to yield SLA and DHAP. Alternatively, a sulfofucolytic ED pathway could involve the oxidation of SFuc to sulfogalactonolactone (SGalL) (by protein MDP0926275), followed by lactone hydrolysis to form SGal (by protein MDP0926279). This intermediate would undergo dehydration to 3-deoxy-2-ketosulfogalactonate (KDSGal) (by protein MDP0926277), and retroaldol cleavage (by protein MDP0926278) to generate *S*-sulfolactaldehyde and pyruvate. In either pathway, *S*-sulfolactaldehyde would be further metabolized by oxidation to *S*-sulfolactate and then the SlcC/ComC–SuyAB biomineralization system encoded in cluster 2 would convert it to sulfite, which would be exported by TauE homologue protein MDP0926575 in cluster 3. Meanwhile DHAP or pyruvate would enter central carbon metabolism.

### Cell-free lysate analysis reveals sulfofucose-stimulated dehydrogenase activity

To explore enzymatic activities in SFuc catabolism, cell-free lysates were prepared from *P. wurundjeri* strain Merri grown in minimal medium with 5 mM SFuc. Cells were harvested in mid-log phase, lysed by sonication, and clarified by centrifugation. NAD(P)^+^-dependent dehydrogenase activity was assessed by monitoring NAD(P)H formation at 340 nm. An increase in absorbance was observed only when lysate and SFuc were present, indicating SFuc-stimulated dehydrogenase activity (**Figure S2**). To test for isomerase activity, ^1^H NMR spectroscopy was used to monitor reactions in lysate supplemented with recombinant shrimp alkaline phosphatase, which prevented phosphorylation and potential downstream turnover of sulfotagatose. After 24 hours of incubation with SFuc, no new resonances consistent with sulfotagatose were detected (data not shown), suggesting the absence of a SFuc–sulfotagatose isomerase. These findings support an ED-like sulfofucolytic pathway in which SFuc undergoes initial NAD(P)^+^-dependent oxidation.

### Detection of sulfogalactonate and sulfolactate by metabolomics analyses

To investigate the metabolic fate of SFuc in *P. wurundjeri* strain Merri, we performed metabolomic analysis on cell lysates from cultures grown with SFuc as the sole carbon source. Using ion chromatography-mass spectrometry (IC-MS), we screened for intermediates indicative of sulfofucolytic EMP- or ED-like pathways. Metabolites were extracted from mid-log phase cell lysates and were analyzed using ion chromatography coupled to high-resolution mass spectrometry.

In SFuc-grown samples, we observed peaks with retention times, *m/z* values and fragmentation patterns matching those of authentic standards for sulfogalactonate (SGal, *m/z* 259) and SL (*m/z* 169) (**Figure 4, Figure S3**). In contrast, no signal corresponding to sulfotagatose, an expected intermediate of an EMP-like pathway, was detected consistent with the lack of SFuc–sulfotagatose isomerase activity in lysate assays. The detection of SGal supports an ED-like pathway, where SFuc is oxidized to sulfofuconolactone, followed by hydrolysis to yield SGal. The presence of SL, the product of SLA oxidation, supports downstream metabolism involving an SLA dehydrogenase, and the SlcC/ComC–SuyAB biomineralization pathway to produce sulfite, detected in the colorimetric assay. Combined with SFuc-stimulated NAD(P)^+^-dependent dehydrogenase activity, this supports the operation of an ED-like sulfofucolytic pathway in *P. wurundjeri*.

**Figure 4.**
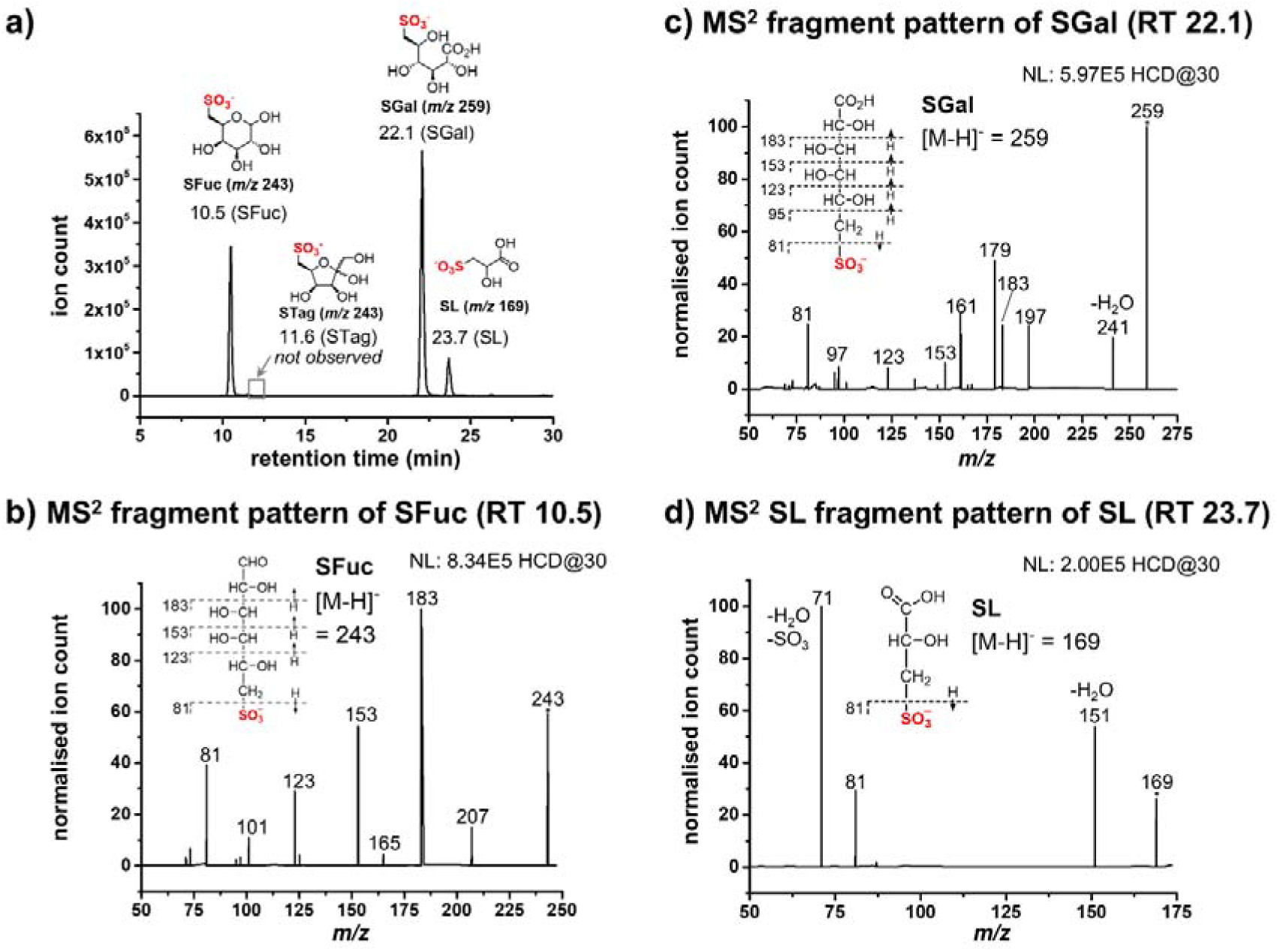
Metabolomic analysis of *P. wurundjeri* strain Merri grown on sulfofucose. Strain Merri cells were grown on SFuc until mid-log phase, harvested, lysed and metabolites were extracted from cells. Targeted LC-IC-MS was employed, and potential sulfofucolytic metabolites were screened for using SIM scans. (a) Extracted ion chromatograph showing key organosulfur metabolites within *P. wurundjeri* cells. (b) MS^2^ fragment pattern of SFuc. (c) MS^2^ fragment pattern of SGal. (d) MS^2^ fragment pattern of SL.

### *In vitro* demonstration of an Enter-Doudoroff pathway for sulfofucose

To demonstrate the biochemical steps of sulfofucolysis, genes encoding the candidate enzymes were heterologously expressed in *E. coli*, and the corresponding proteins were purified. Based on the roles subsequently demonstrated in SFuc catabolism, these genes were renamed *sfcH* (MDP0926275; dehydrogenase), *sfcD* (MDP0926279; lactonase), *sfcF* (MDP0926277 dehydratase), and *sfcE* (MDP0926278; aldolase), reflecting their membership in a <u>s</u>ulfofu<u>c</u>ose degradation pathway. Incubation of SfcH with SFuc and either NADP^+^ or NAD^+^ produced a compound with *m/z* 241, two mass units lower than SFuc, consistent with oxidation to SGalL (**Figure 5**). MS^2^ fragmentation of this species matched that of the diastereoisomer SGL (18). Notably, this enzyme displayed greater activity on NAD^+^, a preference that matches that of SQ dehydrogenase from *P. putida* (37). Subsequent addition of SfcD along with Ca^2+^ and Zn^2+^ converted the oxidized intermediate into a compound with identical retention time and MS^2^ fragmentation pattern as authentic chemically synthesized SGal. When chemically synthesized SGal was incubated with SfcF, a new species with *m/z* 241 (18 mass units lower than sulfogalactonate) was detected, consistent with conversion to KDSGal, and which displayed a fragmentation pattern consistent with the diastereoisomer KDSG (18). Finally, the addition of SfcE and Mg^2+^ to this reaction mixture yielded a product with retention time and MS^2^ fragmentation matching that of chemically synthesized SLA. Separately, analysis of this reaction mixture by IC-MS demonstrated formation of pyruvate (**Figure S4**). SfcE was inactive when supplemented with Mg^2+^ or Mn^2+^.

**Figure 5.**
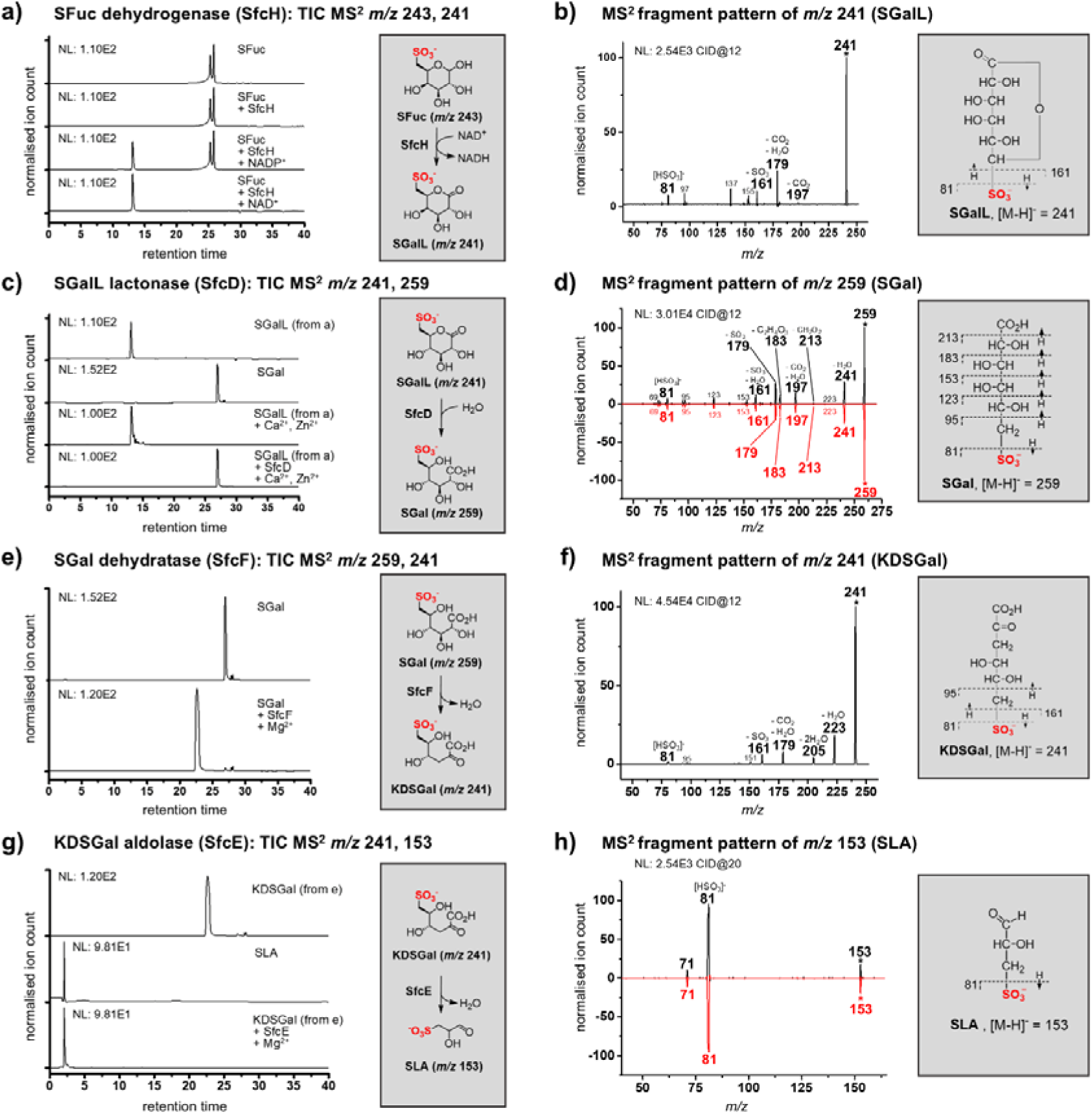
In-vitro reconstitution of individual steps of the sulfofucolytic ED pathway. The enzymatic conversions of SFuc to SGalL, SGalL to SGal, SGal to KDSGal, and KDSGal to SLA, catalyzed by SfcH, SfcD, SfcF and SfcE, respectively. Targeted MRM-MS analysis was used to monitor analytes. (a) Ion chromatograms of SFuc (*m/z* 243) and assays containing SFuc with SfcH and NAD(P)^+^. (b) MS^2^ fragment pattern of SGalL *m/z* 241. (c) Ion chromatograms of assays containing SGalL treated with SfcD, CaCl_2_, and ZnCl_2_. (d) MS^2^ fragment pattern of experimentally obtained SGal (black) and chemically synthesized (red) *m/z* 259. (e) Ion chromatograms of SGal and assays containing SGal with SfcF and MgCl_2_. (f) MS^2^ fragment pattern of KDSGal *m/z* 241. (g) Ion chromatograms of assays containing KDSGal treated with SfcE and MgCl_2_. (h) MS^2^ fragment pattern of experimentally obtained SLA (black) and chemically synthesized (red) *m/z* 153. Pyruvate, the second product of the SfcE reaction, was confirmed by ion chromatography-mass spectrometry (Figure S4).

### Identification of possible sulfofucolytic pathways in proteobacteria

To identify candidate sulfofucolytic ED pathways in other bacteria, we used tools from the Enzyme Function Initiative (EFI) (38, 39) and performed a BLAST search using the sequence of sulfogalactonate (SGal) dehydratase SfcF as a query. Homologous sequences were retrieved and used to construct a sequence similarity network (**Figure 6**). We then examined the genome neighborhoods of these SfcF homologues for the presence of genes encoding additional enzymes associated with the sulfofucolytic ED pathway, specifically, those from PFAM families PF08450 (SGalL lactonase; SfcD), PF13378-PF02746 (SGal dehydratase, SfcF), and PF03328 (KDSGal aldolase, SfcE) (**Table S5**). This analysis identified two clusters of proteins. Gene neighborhood diagrams from the parent bacteria showed that some clusters encode all four core enzymes of the proposed pathway, along with an ABC transporter system. Other clusters lacked an obvious lactonase candidate and some encoded an aldose-1-epimerase, which may function as a SFuc mutarotase. Many of these bacteria also possessed genes encoding homologues of ComC, SlcC, SuyA and SuyB, suggesting that they also possess the capacity for complete metabolism of SFuc through a tandem sulfofucolytic-biomineralization pathway (**Table S6**). Notably, the closest relative of *P. wurundjeri* strain Merri based on 16S rRNA analysis, *P. methylarcula* strain H1, contains syntenic sulfofucolytic and biomineralization gene clusters.

**Figure 6.**
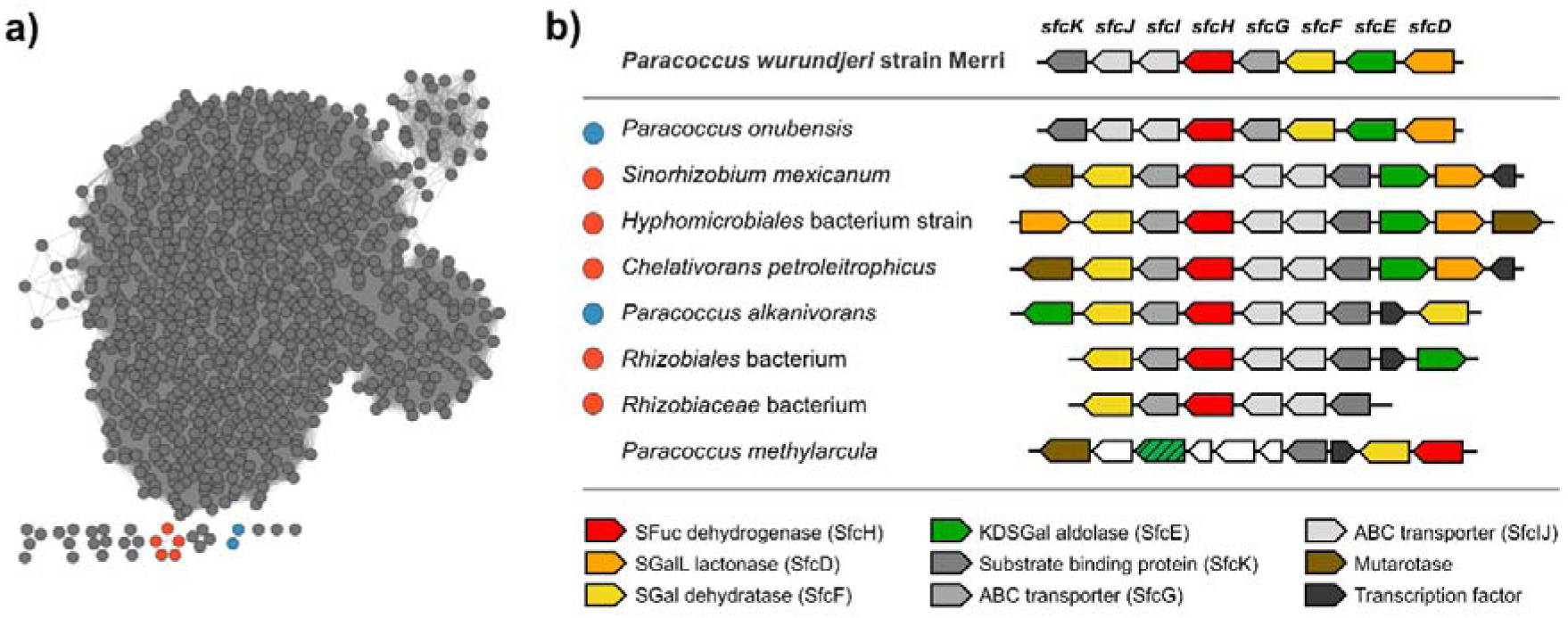
Proposed sulfofucolytic ED pathway gene clusters distributed across assorted proteobacteria. (a) Sequence similarity network (SSN) of SGal dehydratase SfcF homologues at an alignment score of 160. Colored clusters represent organisms harboring gene clusters containing homologues of sulfofucolytic ED pathway genes *sfcE*, *sfcF* and *sfcH*. (b) Gene clusters encoding sulfofucolytic ED pathways. Colored circles at left correspond to colored clusters in part (a). UniProt protein accession codes for SfcF homologues are as follows: *Paracoccus wurundjeri* strain Merri (MDP0926277), *Paracoccus onubensis* strain L29414 (A0A418T7C2), *Sinorhizobium mexicanum* strain ITTG R7 (A0A859QZ22), *Hyphomicrobiales* bacterium strain SCN_18_10_11_15_R4_B_69_13 (A0A8I1P1S0), *Chelativorans petroleitrophicus* strain SCAU2102 (A0A9X2XAH4), *Paracoccus alkanivorans* strain 4-2 (A0A3M0MGN3), *Rhizobiales* bacterium strain 65-79 (A0A1M2Z1S0), *Rhizobiaceae* bacterium strain FW021_bin.52 (A0A522R0Q2), *Paracoccus methylarcula* strain H1 (A0A3R7NEB5).

### A missing step in SLA oxidation to SL

Proteomic and genomic analyses did not identify a homologue of GabD/SlaB capable of oxidising SLA, the product of KDSGal aldolase (SfcE) activity on KDSGal. However, metabolomic analysis revealed the accumulation of SL as an intracellular metabolite during growth of *P. wurundjeri* on SFuc. Within gene cluster 2, comparative proteomics highlighted putative threonine dehydrogenase MDP0926266, which we considered as a candidate SLA dehydrogenase. However, the recombinantly expressed MDP0926266 protein showed no activity toward SLA in the presence of either NAD^+^ or NADP^+^.

## Discussion

In this study, we identify a novel bacterium, *P. wurundjeri sp. nov.*, which can grow on the rare sulfosugar SFuc as sole carbon source and effects its complete degradation to sulfite, via SGal and SL. This organism utilizes a sulfofucolytic Entner–Doudoroff (ED) pathway that cleaves the C6 backbone of SFuc into two C3 fragments: SLA and pyruvate (**Figure 7**). SLA is then biomineralized through a pathway involving oxidation to SL, then sulfolyase catalysed elimination of sulfite to give pyruvate. The pyruvate outputs of this pathway are used in central carbon metabolism, while the sulfite is excreted into the culture media, where it can undergo autooxidation to sulfate. This finding adds to the known metabolic versatility of the genus *Paracoccus*, as *P. pantotrophus* strain NKNCYSA has previously been reported to degrade a range of organosulfonates, including cysteate, taurine, homotaurine, isethionate, and SL (40), suggesting a broad organosulfonate catabolic potential within this taxonomic grouping. However, BLAST analysis reveals that *P. pantotrophus* NKNCYSA does not contain a candidate sulfofucolytic pathway.

**Figure 7.**
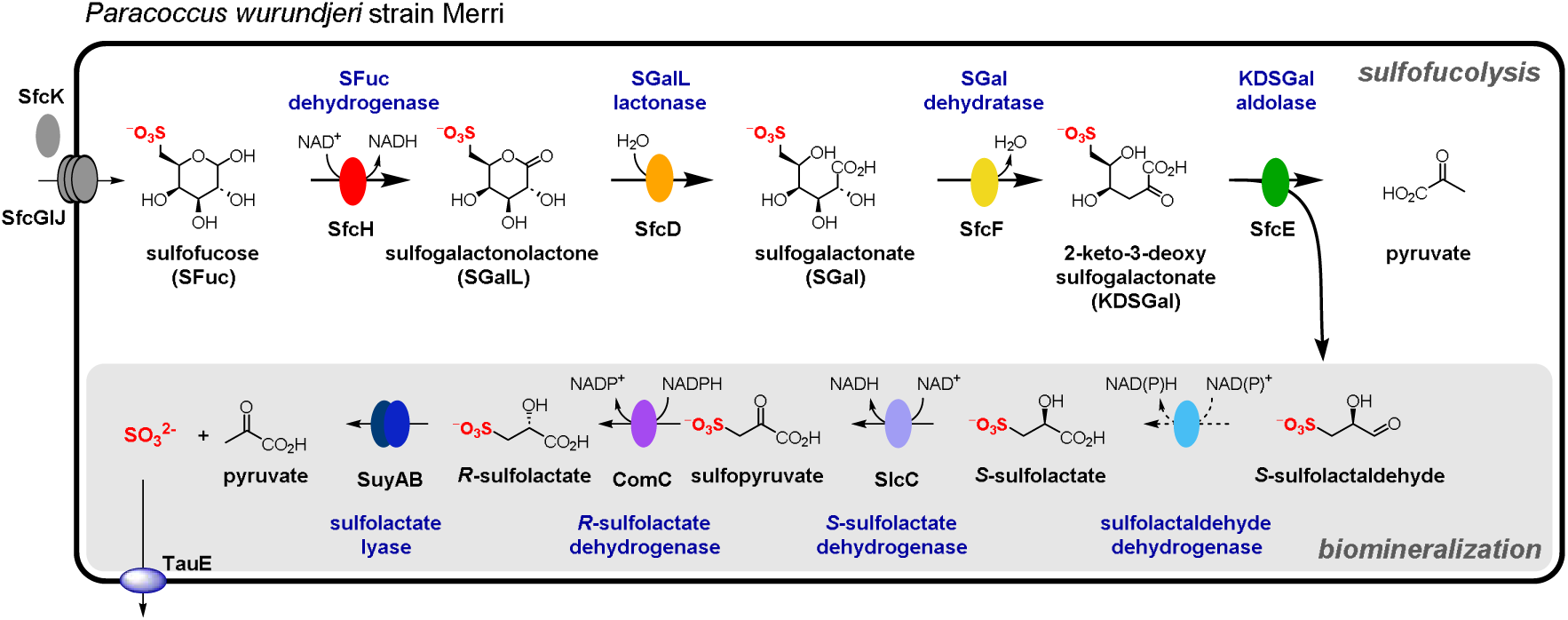
Sulfofucolytic and proposed biomineralization pathway within *P. wurundjeri* strain Merri. The experimentally demonstrated sulfofucolytic steps (bold arrows) involve SFuc dehydrogenase (SfcH), which catalyses the conversion of SFuc to SGalL; SGalL lactonase (SfcD), which catalyses the hydration of SGalL to SGal; SGal dehydratase (SfcF) which catalyses dehydration of SGal to KDSGal; and KDSGal aldolase (SfcE), which catalyses catalyses lysis of KDSGal to yield SLA and pyruvate. The resultant SLA is proposed to undergo a biomineralization process in which *S*-SLA is oxidised to *S*-SL by an unknown SLA dehydrogenase (dotted arrow), racemized by SL dehydrogenases (SlcC/ComC) to *R*-SL, and finally sulfite is liberated by SL lyase (SuyAB) (plain arrows).

The observation of SGal by metabolomic analysis of SFuc-grown *P. wurundjeri* cells is diagnostic of an oxidative sulfofucolytic pathway. We identified four enzymes (SfcH, SfcD, SfcF and SfcE) that collectively enable the stepwise conversion of SFuc through an Entner–Doudoroff–like pathway: SFuc oxidation to SGalL (SfcH), hydrolysis to SGal (SfcD), dehydration to KDSGal (SfcF), and aldol cleavage to yield SLA and pyruvate (SfcE). These steps mirror the enzymatic steps of the canonical sulfoglycolytic ED pathway used for SQ catabolism, with the key distinction to SFuc being the stereochemistry at C4 of the sugar. This renders their corresponding intermediates epimeric: SGalL vs. SGL, SGal vs. SG, and KDSGal vs. KDSG. Despite this difference, the aldolase-catalyzed cleavage of both KDSG and KDSGal generates the same *S*-enantiomer of SLA. This is because the stereocenter in SLA originates from carbon 5 which is common to both sulfosugars, while the configuration at carbon 4 is lost during cleavage, and so does not affect the stereochemistry of the product.

All but one of the enzymes in the sulfofucolytic ED pathway belong to the same protein families as those in the canonical sulfoglycolytic ED pathway for sulfoquinovose catabolism (18). The key difference lies in the dehydratase step: while sulfoquinovose degradation employs SG dehydratase (PF00920), the sulfofucolytic pathway utilizes SGal dehydratase, which belongs to a distinct pair of families (PF13378, PF02746). This divergence provided a basis for a targeted bioinformatic search, enabling the identification of related gene clusters in other bacteria. Using this signature, we identified a small subset of Proteobacteria harboring syntenic gene clusters that encode homologues of the sulfofucolytic ED pathway. Notably, like that of strain Merri, all of these gene clusters were associated with ABC transporter systems, which may mediate SFuc uptake. ABC transporters have previously been invoked in SQ uptake in sulfo-ED pathways (18, 41), and SQ-binding proteins associated with ABC transporters have been characterized biochemically and structurally (27, 42, 43). Several organisms contained a candidate SFuc mutarotase, which could facilitate the oxidation of SFuc to SGalL if the SFuc dehydrogenase displays specificity for one of the anomers of SFuc by accelerating the rate of anomer interconversion. Notably, anomeric preference has been reported in the first step of the sulfo-EMP pathway in *E. coli*, where SQ isomerase preferentially acts on the β-anomer of SQ during its isomerization to sulfofructose (44).

Interestingly, several of the candidate sulfofucolytic organisms lacked an obvious lactonase homologue, raising the possibility that they utilize a variant of the pathway that bypasses the lactone hydrolysis step. This difference is reminiscent of the De Ley–Doudoroff pathway—a variant of the Entner–Doudoroff pathway—in which D-galactose is oxidized by a NAD⁺-dependent dehydrogenase to D-galactonate, followed by dehydration to 2-keto-3-deoxygalactonate and subsequent retro-aldol cleavage to yield pyruvate and glyceraldehyde (45, 46). By analogy, these bacteria may encode a lactonase-independent sulfofucolytic ED pathway in which SFuc is directly oxidized to SGal, thereby bypassing the need for SGalL hydrolysis.

Comparative proteomics of strain Merri identified a cluster of genes that encoded homologues of proteins capable of the biomineralization of the C3-organosulfonate SL. Specifically, these genes encode homologues of the two component NAD(P)^+^-dependent SL racemases SlcC/ComC (25), and SL lyase (SuyAB), which cleaves SL to yield pyruvate and sulfite (35, 36). While no homologues of classical SLA dehydrogenases SlaB/GabD (18, 19, 47) were present, a candidate SLA dehydrogenase (MDP0926266) was identified that belonged to the protein families PF08240-PF00107 and was annotated as threonine dehydrogenase; however, recombinant MDP0926266 did not oxidize SLA. Nonetheless, metabolomic analysis of lysates from SFuc-grown cells revealed the presence of SL, consistent with the existence of this step in the pathway. Together, these data provide support for pathway involving SLA dehydrogenase and the SlcC/ComC–SuyAB biomineralization pathway, proceeding via the sequence *S*-SLA → *S*-SL → sulfopyruvate →*R*-SL → pyruvate + sulfite. Genes encoding homologues of SlcC, ComC, SuyA and SuyB were present in several of the bacterial strains encoding candidate sulfofucolytic pathways, namely, *Paracoccus alkanivorans*, *Rhizobiaceae* sp., *Sinorhizobium mexicanum*, *Chelativorans petroleitrophicus*, and *Paracoccus onubensis*, suggesting that these bacteria also utilize tandem sulfofucolytic-biomineralization pathways (**Table S6**).

This work provides evidence for the coexistence of both sulfofucolysis and biomineralization pathways within a single bacterium. Co-existence of a sulfofucolytic/sulfoglycolytic and biomineralization pathway in a single bacterium is unusual. One example involves an environmental *Klebsiella* sp., which was shown to degrade SQ to SL and, after 24 h growth, produced trace amounts of sulfate—equivalent to approximately 10% of the initial SQ (48). However, the molecular mechanisms underlying these observations in that *Klebsiella* strain remain unresolved.

The occurrence of SFuc has only been documented in two reports: within glycoconjugates produced by *Fucacaea* algae and an extremophile archaeon (28, 29). This limited evidence suggests that SFuc may be relatively rare in the environment. However, members of the *Fucacaea* are widespread in coastal regions around the North Altantic Ocean, raising the possibility that SFuc is more common than currently recognised. Moreover, SFuc shares an identical molecular formula with sulfoquinovose (SQ), meaning it could have been misidentified in studies that relied exclusively upon mass spectrometry for identification. If so, its true environmental distribution may be broader than literature reports suggest.

Strain Merri was isolated from Joe’s Garden, a riparian market garden located on the banks of Merri Creek in Melbourne, Australia. Given the site’s history of flooding, the bacterium, or its substrate, may have a soil or aquatic origin. The Proteobacteria harboring syntenic sulfofucolytic gene clusters have been isolated from soil and oil-rich environments, and include rhizobia, indicating that SFuc may be encountered in soil ecosystems, particularly within the rhizosphere. In these niches, bacteria may encounter SFuc released from known or yet-unidentified SFuc-containing glycoconjugates. Although no candidate sulfofucosidases were identified in this study, previous work has shown that sulfoquinovosidases from *E. coli* and *Agrobacterium tumefaciens* are inactive on PNP α-sulfofucoside (15), suggesting limited enzymatic cross-reactivity and the potential for discovery of a novel sulfofucosidase. Finally, we highlight the existence of another sulfohexose, sulforhamnose, which arises transiently as an intermediate in the sulfo-EMP pathway from SQ and is subsequently isomerized back to SQ or sulfofructose, key intermediates in that pathway (44).

In conclusion, this work identifies a bacterium, *P. wurundjeri*, capable of complete catabolism of the rare sulfosugar SFuc. We provide evidence for a two-part metabolic route: an oxidative sulfofucolytic Entner–Doudoroff (ED) pathway that cleaves SFuc into C3-organosulfonate intermediates, and a downstream biomineralization pathway involving SL lyase–mediated conversion of SL to sulfite. While most enzymes in the sulfofucolytic ED pathway share homology with those in the known sulfoglycolytic ED pathway, the dehydratase step is catalyzed by a protein from a distinct family (PF13378, PF02746). This divergence enabled the identification of a small set of bacteria predicted to possess similar SFuc catabolic pathways. Notably, *P. wurundjeri* did not grow on SFuc-containing glycosides, indicating the absence of a sulfofucosidase capable of liberating free SFuc from glycoconjugates (data not shown). Future studies could focus on identifying and characterizing such sulfofucosidases, which would expand our understanding of how SFuc is accessed and degraded in diverse environmental contexts.

## Materials and Methods

### Specialist chemicals

Sulfolactate and sulfolactaldehyde were synthesized as reported (49, 50). The synthesis of sulfofucose and sulfogalactonate from D-galactose, ^13^C_6_-sulfofucose from ^13^C_6_-D-galactose, and sulfotagatose from D-galacturonic acid will be reported elsewhere.

### Bacterial growth media

Growth media was prepared using M9 minimal salts media (2 ml), trace metal solution (0.1 ml), and vitamin solution (0.01 ml) and contained 10 mM SFuc as sole carbon source, made up to a final volume of 10 ml with deionized water. M9 minimal salts media contains 0.45 M Na_2_HPO_4_, 0.11 M KH_2_PO_4_, 0.09 M NH_4_Cl, 0.04 M NaCl, 0.1 M MgSO_4_, 0.1 M CaCl_2_. Trace metal solution contains 0.4 mM FeCl_3_, 0.08 mM CoCl_2_, 0.08 mM CuCl_2_, 0.08 mM MnCl_2_, 0.55 mM ZnCl_2_, 0.01 mM NiCl_2_, 0.05 mM Na_2_MoO_4_ and 0.5 mM H_3_BO_3_. Vitamin mixture contains 0.04 mM biotin, 0.05 mM calcium pantothenate, 0.15 mM thiamine hydrochloride, 0.36 mM p-aminobenzoic acid, 0.81 mM nicotinic acid, 1.49 mM pyridoxamine dihydrochloride, 0.01 B12 (cyanocobalamin). Growth curves were measured using a custom built MicrobeMeter portable spectrophotometer.(51)

### Isolation of *Paracoccus wurundjeri* strain Merri

*P. wurundjeri* strain Merri was isolated by substrate-guided enrichment culturing, from soil sampled at Joe’s Market Garden (34 Edna Grove, Coburg, Victoria 3058; 37.7451° S, 144.9813° E). Approximately 1 g of soil was suspended in 5 ml of sterilized M9 media (supplemented with trace metals and vitamins) containing 10 mM SFuc as sole carbon source. The culture was incubated at 30 °C for 4 d with agitation at 250 rpm. A subsample (100 µL) was transferred into fresh M9 media (supplemented with trace metals and vitamins) and grown for a further 4 d. This step was repeated five times and after outgrowth of the final culture for 4 d, cells were plated onto SFuc-agar plates (20 mM SFuc, 15 g/l agar) and incubated at 30 °C in the dark. After 3 d, a single colony was picked and inoculated into fresh M9 media (supplemented with trace metals and vitamins) containing 10 mM SFuc and incubated at 30 °C while shaking at 250 rpm. Once the culture was visibly turbid, cells were again plated onto SFuc-agar plates, incubated at 30 °C in dark and single colony picked and inoculated again into fresh vitamin supplemented M9 media containing 10 mM SFuc. Once the culture was visibly turbid, frozen stocks were prepared by diluting to 5% DMSO and freezing at -80 °C. This strain could grow on racemic DHPS (5 mM), but did not grow on SQ (5 mM), or *S/R*-SL (5 mM).

### Genome sequence, assembly and annotation

*P. wurundjeri* strain Merri was grown in 10 ml M9 media supplemented with 10 mM SFuc for 5 days and then the cell pellet was harvested by centrifugation. Genomic DNA was isolated using the GenElute DNA extraction kit (Sigma) with inclusion of lysozyme and RNAase, then it was sequenced using Illumina NextSeq platform (Doherty Microbial Genomics, University of Melbourne, Australia). DNA was prepared for sequencing using the Nextera XT DNA preparation kit (Illumina) with ×150 bp paired end chemistry and with a targeted sequencing depth of >50×. A draft genome was assembled using Shovill v1.0.9 (https://github.com/tseemann/shovill) and annotated using PGAP as part of the NCBI Genome submission pipeline. GTDB-TK was used to determine average nucleotide identity and taxonomic placement (52–54).

### 13C-NMR analysis of spent culture media from growth of *P. wurundjeri* strain Merri on (^13^C_6_)-sulfofucose

M9 minimal media (supplemented with trace metals and vitamins) (4 ml) containing 10 mM (^13^C_6_)-SFuc was inoculated with *P. wurundjeri* strain Merri and grown to mid-log phase (5 days) and stationary phase (14 days) at 30 °C (250 rpm). The final A_600_ was 0.3 (5 days) and 0.52 (14 days). Cells were sedimented by centrifugation at 5,000 × *g* for 3 min, and the supernatant was collected. 540 μL of culture supernatant was diluted with 60 μL D_2_O and ^13^C-NMR spectra acquired using a 500 MHz spectrometer. A control sample contained 10 mM (^13^C_6_)-SFuc in M9 minimal media prepared in the same way for NMR analysis.

### Analysis of sulfofucose, sulfite and sulfate in *P. wurundjeri* culture

M9 minimal media (supplemented with trace metals and vitamins) 50 mL containing 10 mM SFuc was inoculated with *P. wurundjeri* strain Merri and grown at 30 °C at 250 rpm in a 500 mL baffled conical flask. Samples were withdrawn at various time points, cells were pelleted by centrifugation for 5 min at 5,000 *g*, and supernatant samples were frozen and stored at -80 °C prior to analysis for reducing sugar, sulfate, or sulfite content.

### Reducing sugar assay

The reducing sugar assay was performed according to the procedure of Blakeney and Mutton (55). Alkaline diluent was prepared by the addition of sodium hydroxide (20 g, 0.50 mol) to a solution of 0.10 M trisodium citrate (50 mmol, 500 ml) and 0.02 M calcium chloride (13 mmol, 500 ml). PAHBAH working solution was prepared by dissolving 4-hydroxybenzhydrazide (PAHBAH) (0.25 g, 1.6 mmol) in alkaline diluent (50 ml). The PAHBAH working solution should be made fresh shortly before use. To determine reducing sugar concentration, 90 µL of PAHBAH working solution and 10 µL of culture supernatant were combined in a 96-well half-area microplate. The mixture was heated at 98°C for 5 min, and the absorbance was read at 415 nm using a UV/visible spectrophotometer. Concentrations of SFuc were determined by reference to a standard curve constructed using SFuc. Analyses were performed in duplicate and errors are reported as standard deviation.

### Turbidometric sulfate assay

The sulfate assay was performed according to the procedure of Sörbo.(56) This assay uses a barium sulfate–polyethylene glycol (Ba-PEG) reagent, which contains polyethylene glycol (PEG) to stabilize BaSO_4_ crystals and a small amount of preformed BaSO_4_ seed crystals to improve the reproducibility and linearity of the assay. Ba-PEG reagent was prepared by dissolving BaCl_2_ (42 mg, 0.20 mmol) and polyethylene glycol 6000 (0.75 g) in deionized water (5.0 ml). A small amount of Na_2_SO_4_ (10 μl, 50 mM) was added to this solution, with efficient magnetic stirring to generate preformed BaSO_4_ seed crystals. The Ba-PEG reagent must be prepared fresh on the day of use. Individual sulfate assays were conducted as follows: In an Eppendorf tube, 50 μl of 0.5 M HCl was added to 50 μl culture supernatant, followed by Ba-PEG reagent (50 μl). The mixture was mixed vigorously and then 90 μl was transferred to a 96-well half-area microplate. The absorbance of the sample at 400 nm was measured using a UV/visible spectrophotometer. Concentrations of sulfate were determined by reference to a standard curve constructed using 0-10 mM Na_2_SO_4_. Analyses were performed in duplicate and errors are reported as standard deviation.

### Colorimetric fuchsin sulfite assay

The fuchsin sulfite assay was performed according to the procedures of Brychkova et al.(57) and Kurmanbayeva et al.(58) This method requires three freshly pre-prepared solutions: Reagents A, B, and C. Reagent A was prepared by adding 1 ml Schiff’s reagent to 8 ml deionised water, followed by addition of 5 M HCl until the solution turned colorless. Reagent B was prepared by diluting formaldehyde (36% in H_2_O, 0.32 ml) in deionised water (9.68 ml) at 0 °C. Reagent C was prepared by dilution of Reagent A (1 ml) in deionised water (7 ml), prior to the addition of solution reagent B (1 ml). Individual sulfite assays were performed by adding Reagent C (100 μl) to the cell supernatant (10 μl). The sample was incubated at 25 °C for 10 min and then transferred to a 96 well half area microplate. The absorbance of the sample at 570 nm was determined by using a UV/visible spectrophotometer. Concentrations of sulfite were determined by reference to a standard curve constructed using Na_2_SO_3_. Analyses were performed in duplicate and errors are reported as standard deviation.

### Comparative proteomics

Six independent cultures of *P. wurundjeri* strain Merri were grown in 10 mM SFuc and another six independent cultures were grown in 10 mM glucose (both in M9 media supplemented with trace metals and vitamins). The cultures were harvested at mid-log phase (A_600_ approx 0.25), and bacteria were collected by centrifugation at 4000 × *g* for 10 min. The pelleted cells were washed by resuspension with fresh 10 ml ice-cold phosphate-buffered saline (PBS) and re-pelleting three times, with the supernatant discarded each time. Cells were resuspended and transferred to a 1.5 ml tube using ice-cold PBS (1 ml), and collected by centrifugation for 2 min at 10,000 × *g*, and then snap-frozen in liquid nitrogen.

Frozen whole bacterial pellets were prepared using the in-StageTip preparation approach as previously described.(59) Cells were resuspended in 4% sodium deoxycholate (SDC), 100 mM Tris pH 8.0 and boiled at 95 °C with shaking for 10 min to aid solubilisation. Samples were allowed to cool for 10 min and then boiled for a further 10 min before the protein concentration was determined by bicinchoninic acid assays (Thermo Fisher Scientific). 100 μg of protein equivalent for each sample was reduced/alkylated with the addition of tris-2-carboxyethylphosphine hydrochloride and iodoacetamide (final concentrations 20 mM and 60 mM, respectively), by incubating in the dark for 1 h at 45 °C. Following reduction/alkylation, samples were digested overnight with trypsin (1/33 w/w Solu-trypsin, Sigma) at 37 °C with shaking at 1000 rpm. Digests were then halted the addition of isopropanol and trifluoroacetic acid (TFA) (50% and 1%, respectively) and samples were cleaned up using home-made SDB-RPS StageTips prepared according to previously described protocols.(59–61) SDB-RPS StageTips were placed in a Spin96 tip holder 2 to enable batch-based spinning of samples and tips conditioned with 100% acetonitrile; followed by 30% methanol, 1% TFA followed by 90% isopropanol, 1% TFA with each wash spun through the column at 1000 *g* for 3 min. Acidified isopropanol / peptide mixtures were loaded onto the SDB-RPS columns and spun through, and tips were washed with 90% isopropanol, 1% TFA followed by 1% TFA in deionised water. Peptide samples were eluted with 80% acetonitrile, 5% ammonium hydroxide and dried by vacuum centrifugation at room temperature, then were stored at -20 °C.

Prepared digested proteome samples were re-suspended in Buffer A* (2% acetonitrile, 0.01% TFA) and separated using a two-column chromatography setup composed of a PepMap100 C18 20-mm by 75-μm trap and a PepMap C18 500-mm by 75-μm analytical column (Thermo Fisher Scientific). Samples were concentrated onto the trap column at 5 μl/min for 5 min with Buffer A (0.1% formic acid, 2% DMSO) and then infused into an Orbitrap Fusion Eclipse at 300 nl/minute via the analytical columns using a Dionex Ultimate 3000 UPLCs (Thermo Fisher Scientific). 125-minute analytical runs were undertaken by altering the buffer composition from 2% Buffer B (0.1% formic acid, 77.9% acetonitrile, 2% DMSO) to 22% B over 95 min, then from 22% B to 40% B over 10 min, then from 40% B to 80% B over 5 min. The composition was held at 80% B for 5 min, and then dropped to 2% B over 2 min then was held at 2% B for another 8 min. The Fusion Eclipse Mass Spectrometer was operated in a data-dependent mode with a single Orbitrap MS scan (375-1500 *m/z* and a resolution of 120k) was acquired every 3 seconds followed by Orbitrap MS/MS higher energy collisional dissociation (HCD) scans of precursors (stepped NCE 25,30,40%, with a maximal injection time of 80 ms with the Automatic Gain Control set to 400% and the resolution to 30k).

Identification and LFQ analysis were accomplished using MaxQuant (v1.6.17.0)(62) using the in-house generated proteome of *P. wurundjeri* strain Merri (RES21-00833.faa) allowing for oxidation on methionine. The LFQ and “Match Between Run” options were enabled to allow comparison between samples. The resulting data files were processed using Perseus (v1.4.0.6)(63) to compare the growth conditions using Student’s t-tests as well as Pearson correlation analyses. For LFQ comparisons independent replicates were grouped and missing values imputed based on the observed total peptide intensities with a range of 0.3σ and a downshift of 1.8σ.

### Cell free lysate analysis

*P. wurundjeri* strain Merri cells were grown in vitamin-supplemented M9 minimal media (250 mL) containing 5 mM SFuc. Cell pellets were collected at mid log phase by centrifugation at 4000 × *g* for 10 min at 5 °C washed thrice with cold PBS buffer (150 mM), thrice with cold water, and frozen on liquid nitrogen. Cells were defrosted and resuspended in 100 mM Tris-HCl buffer (pH 7.0, 100 mM) and lysed by sonication with 30% powder in 3 x 30 sec intervals. Samples were cooled on ice for 1 minute in-between cycles. Samples were clarified by centrifuge at 25,000 × *g* for 10 min at 5 °C and the cell lysate was used without further treatment.

### Sulfofucose isomerase assay

Potential SFuc isomerase reactions were monitored by ^1^H NMR spectroscopy using a 500 MHz Bruker NEO system. A 700 μL sample containing 50 μL of strain Merri cell lysate, 50 units of recombinant shrimp alkaline phosphatase (Sigma), in 50 mM sodium phosphate, 150 mM NaCl (pH 7.01) was incubated for 20 min. SFuc was added to a final concentration of 10 mM and samples were incubated for 24 h at 25 °C. Reactions were quenched by heating at 80 °C for 3 min, concentrated *in vacuo* and the residue was dissolved in D_2_O for ^1^H NMR analysis.

### Sulfofucose dehydrogenase assay

In a 96-well plate, 200 μL solutions contained 1.1 mM SFuc, 6.5 mM MgCl_2_, and 0.5 mM NAD^+^ or NADP^+^ in Tris-HCl (100 mM, pH 7.5). Reactions were initiated by the addition of 20 μL of strain Merri cell lysate. Absorbance was measured at 340 nm for 15 mins.

### Metabolomic analysis of *P. wurundjeri* strain Merri

Three independent replicates of *P. wurundjeri* strain Merri were grown to mid log phase in vitamin-supplemented M9 minimal media with either 5 mM SFuc or glucose as the sole carbon source. Samples were grown to the following A_600_ values: SFuc: 1.126, 1.130, 1.123 Glucose: 1.06, 1.01, 1.10. Cultures were centrifuged (4000 rpm, 30 cm rotor, 10 min, 5 °C) and cell pellets were washed three times with PBS buffer (10 mM, NaCl 150 mM, pH 7.0) and then three times with deionised water. Cells were snap frozen with liquid nitrogen prior to extraction.

250 µL of MTBE:MeOH (3:1 v/v) and 125 µL of H_2_O/MeOH (3:1 v/v) were added to the frozen cell pellets and cells were lysed by sonication in a soniclean 160TD at 100% power for 10 min. Solutions were then mixed at 4 ° C for 30 min at 1200 rpm in a ThermoMixer. Samples were centrifuged at 6,000 × *g* for 45 min and the aqueous phase was separated and evaporated to dryness. The residues were reconstituted with 40µL of MeOH:H_2_O (2:3 v/v) and the metabolome of *P. wurundjeri* strain Merri was analysed by ion chromatography-mass spectrometry (IC-MS).

IC-MS analysis was conducted on a ThermoFisher Scientific LC ID-X Orbitrap equipped with a Dionex IonPac AS11-HC column (ThermoFisher Scientific).

A 10 µL sample aliquot was introduced onto a heated (30 °C) Dionex™ IonPac™ AS11-HC analytical column (2 mm × 250 mm, 9 μm particle size) with its corresponding AG11-HC guard column (2 mm × 50 mm) using a Dionex™ ICS-6000 HPIC system (Thermo Scientific, USA) equipped with an integrated KOH eluent generator using a flow rate of 380 µL/min. The elution gradient consisted of 2 mM KOH held for 0.3 min, followed by linear increases to 12 mM over 13.2 min, to 20 mM over 9 min, and to 70 mM over an additional 9 min. The concentration was maintained at 70 mM for 6 min before returning to 2 mM within 0.1 min, with a subsequent re-equilibration period of 9.4 min. Sample diversion to the mass spectrometer began at the 2-minute mark. An AXP-MS pump (Thermo Scientific, USA) delivered acetonitrile as make-up flow at 0.12 mL/min, which was introduced into the eluent stream via a T-split (Thermo Scientific, USA) positioned downstream of the conductivity detector and as proximal to the ionization source as feasible.

MS acquisition was performed using a Thermo Scientific ID-X system operating at 240,000 resolution (referenced at m/z 200). The chromatographic eluent was introduced through a heated electrospray ionization (HESI) source maintained at 225 °C with optimized gas parameters (sheath: 58 arbitrary units, auxiliary: 14 arbitrary units, sweep: 0.5 arbitrary units). Metabolites were ionized in negative mode using a 3000 V spray voltage with mild trapping enabled and RunStartEasyIC™ calibration with selected ion monitoring in negative ionization mode to monitor for SFuc [M-H]^-^ *m/z* 243, SGalL [M-H]^-^ *m/z* 241, SGal[M-H]^-^ *m/z* 249, KDSGal [M-H]^-^ *m/z* 241, SLA [M-H]^-^ *m/z* 153 and SL [M-H]^-^ *m/z* 169. Ions monitored were fragmented by CID (0-30 arbitrary units) or HCD (0-45 arbitary units). Additional desolvation occurred in the transfer tube heated to 300 °C. Data were collected across a mass range of 100-1000 m/z with default maximum injection time and automatic gain control (AGC) target settings, and an RF lens value of 35%. Fragmentation spectra were acquired using data-dependent acquisition (ddMSn) with higher energy collisional dissociation (HCD) at assisted stepped collision energies (15, 30, and 45%) at 60,000 resolution using a 1 Dalton isolation window. A second ddMSn event employed collision-induced dissociation (CID) at 30% collision energy with a 1.6 Dalton isolation window at the same resolution. All scans used [M-H]^-^ *m/z* 243, 241, 259, 169, 153 inclusion lists.

### Cloning, protein expression and purification of SfcD, SfcE, SfcF, SfcH, and MDP0926266

The codon-harmonized protein-coding regions of SfcD, SfcE, SfcF, SfcH, and MDP0926266 genes from *P. wurundjeri* strain Merri (MDP0926279, MDP0926278, MDP092627, MDP0926275, and MDP0926266, respectively) were synthesized and cloned between *Nco*I and *Xho*I in pET-28a with an N-terminal hexahistidine tag, followed by the TEV protease recognition site. For protein production, plasmids encoding SfcE, SfcF, SfcG, or MDP0926266 was transformed in *E. coli* BL21(DE3) and expressed. The cells were grown in lysogeny broth with 50 mg/ml kanamycin at 37 °C and when they reached an A_600_ of 0.6–0.8 were induced with 0.4 mM isopropyl 1-thio-β-D-galactopyranoside (IPTG). After incubation at 18°C for 16 h, cells were harvested by centrifugation at 6000 × *g*. Due to low solubility of the SfcD protein, the plasmid encoding SfcD was expressed in ArcticExpress (DE3) (Agilent). The transformed cells were grown in LB media at 30 °C and when they reached an A_600_ of 0.6–0.8 were induced with 0.1 mM IPTG and incubated at 11 °C for 24 h then harvested as above. The cell pellets were resuspended in buffer A (20 mM Tris-HCl, 500 mM NaCl, 25 mM imidazole, pH 7.5), disrupted using a sonicator (Omni Sonic Ruptor 400), and clarified by centrifugation at 20000 × *g*. The soluble fraction was applied to Ni Sepharose 6 Fast Flow (Cytiva) resin and the bound protein was eluted with 300 mM imidazole in buffer A. Pooled fractions were concentrated and buffer exchanged into 20 mM Tris-HCl, 200 mM NaCl, pH 7.5.

### Analysis of the products of reactions catalysed by SfcD, SfcE, SfcF and SfcH

SFuc dehydrogenase (SfcH): A 0.5 mL solution contained SFuc (2 mM), and either NAD^+^ (3 mM) or NADP^+^ (3 mM) in Tris-HCl buffer (50 mM, pH 8.0). SfcG (1.67 µM) was added to initiate the reaction and the solution was incubated at 25 °C for 24 h, then halted by heating at 80 °C for 3 min. Samples were frozen until LC-MS analysis.

SGalL lactonase (SfcD): A 0.3 mL solution contained 100 µL of the final SfcG solution from above, CaCl_₂_ (0.5 mM), and ZnCl_₂_ (0.5 mM) in Tris-HCl buffer (50 mM, pH 8.0). The reaction was initiated by adding SfcG (2.67 µM) and the solution was incubated at 25 °C for 24 h, then halted by heating at 80 °C for 3 min. Samples were frozen until LC-MS analysis.

SGal dehydratase (SfcF): A 0.5 mL solution contained chemically-synthesized SGal (2 mM) and MgCl_₂_ (1 mM) in Tris-HCl buffer (50 mM, pH 8.0). The reaction was initiated by adding SfcF (1.11 µM) and incubated at 25 °C for 24 h, then halted by heating at 80 °C for 3 min. Samples were frozen until LC-MS analysis.

KDSGal aldolase (SfcE): A 0.5 mL solution contained 100 µL of the final SfcF solution from above, and either MgCl_₂_ (1 mM), MnCl_₂_ (1 mM), or ZnCl_₂_ (1 mM) in Tris-HCl buffer (50 mM, pH 8.0). The reaction was initiated by adding SfcE (1.68 µM) and incubated at 25 °C for 24 h, then halted by heating at 80 °C for 3 min. Samples were frozen until LC-MS analysis.

LC-MS analysis: 20 µL of each sample was diluted with 60 µL of 1:1 ACN/H_2_O and used for LC-MS analysis. An Aglient 6490 QQQ mass spectrometer equipped with an Aglient LC module was used for HPLC–MS analysis. A SeQuant ZIC-HILIC 3.5 µm 100 Å 150 x 2.1 mm column was used at room temperature. HPLC used solvent A, 10 mM NH_4_OAc in 10% acetonitrile; solvent B: acetonitrile. The elution gradients were from 90% B to 40% B over 50 min; then 40% B for 2 min; back to 90% B over 1 min with flow rate of 0.50 ml min^-1^ and injection volume of 1 µL. The mass spectrometer was operated in negative ionization mode. Prior to analysis, the sensitivity for each multiple reaction monitoring (MRM)-MS^2^ transition was optimized for each analyte.

Retention times and MS^2^ fragmentation patterns of the authentic reference standards were as follows:

For SFuc: Retention time: 25.3 and 25.8 min. ESI–MS/MS *m/z* of [M-H]^-^ 243, product ions 283, 153, 123, 81.

For SGalL: Retention time: 13.1 min. ESI–MS/MS *m/z* of [M-H]^-^ 241, product ions 197, 179, 161, 81.

For SGal: Retention time: 27.0 min. ESI–MS/MS *m/z* of [M-H]^-^ 259, product ions 241, 213, 197, 183, 179, 161, 81.

For KDSGal: Retention time: 22.6 min. ESI–MS/MS *m/z* of [M-H]^-^ 241, product ions 223, 205, 179, 161, 81.

For SLA: Retention time: 2.1 min. ESI–MS/MS *m/z* of [M-H]^-^ 153, product ions 81, 71.

Ion chromatography-Mass spectrometry (IC-MS) analysis of pyruvate: Following incubation of KDSGal aldolase reaction mixtures, the enzyme was precipitated by adding CHCl□ (300 µL), then centrifuged at 1400 rpm (using a 15 cm rotor) for 10 minutes. The aqueous layer was collected and diluted to a final pyruvate concentration of 10 µM for IC-MS analysis in a manner analogous to that previously described with selected ion monitoring in negative ionization mode to monitor for pyruvate [M-H]^-^ *m/z* 87. All scans used [M-H]^-^ *m/z* 87 inclusion lists.

### Investigation of activity of MDP0926266

Solutions (150 μL) were prepared containing 1.0 mM racemic SLA, 1.5 mM NAD^+^ or NADP^+^ in 100 mM Tris-HCl (pH 7.5), with or without 0.5 mM ZnCl_2_, FeCl_2_ or FeCl_3_. Reactions were initiated by the addition of MDP0926266 to a final concentration of 30 nM. Absorbance at 340 nm was monitored over 15 minutes. No enzymatic activity was detected under any of the tested conditions.

### Bioinformatic analysis

The protein sequence of SfcF (UniProt accession code: Q5LUS2) from *P. wurundjeri* strain Merri was used as a query for BLAST search in the Enzyme Function Initiative-Enzyme Similarity Tool (EFI-EST webtool, ENA database, version June 2022; UniProt release 2022_02 and InterPro 89) (38, 39). This search retrieved 1001 sequences with ≥ 62.2% identity. Sequence similarity networks (SSNs) were generated at different alignment scores (AS = 120, 140, 160, 180, 200) and visualised in Cytoscape (64) 3.10.3 using the yFiles Organic Layout, with nodes representing individual sequences.

Genome neighborhoods were analyzed using the Enzyme Function Initiative-Genome Neighborhood Tool (EFI-GNT), with nodes color-coded according to colocation with SfcD, SfcE, SfcF, SfcH homologues. Analysis at AS = 160 identified two clusters containing 7 sequences with SfcF and at least one homologue of SfcD, SfcE or SfcH.

Further analysis of the SSN at AS = 160 using EFI-GNT retrieved neighboring genes within a ±10 open reading frame window.

## Supporting information

Supplementary Gene and Protein Sequences

Supplementary Proteomics

## Data Availability

The assembled genome has been deposited at the NCBI (GenBank accession: Bioproject PRJNA993841), release date Aug 7, 2023.

The mass spectrometry proteomics data has been deposited in the Proteome Xchange Consortium(65) via the PRIDE partner repository (https://proteomecentral.proteomexchange.org/cgi/GetDataset) with the data set identifier: PXD043488 Username: Password: DXxRoG1Y.

## Supplemental material

Supplementary Figures S1-S8

Supplemental Tables S1 to S6

Supplemental Information Gene sequences

Supplemental Proteomics data.xlsx

## Conflict of interest

The authors declare that they have no conflict of interest.

## Ethics approval

Not applicable.

## Consent to participate

Not applicable.

## Acknowledgements

This work was supported by the Australian Research Council (DP210100233, DP210100235, DP210100362, DP230102668, DP240100126; DP250100819) and National Health and Medical Research Council of Australia (NHMRC) (GNT2021638). N.E.S is supported by an Australian Research Council Future Fellowship (FT200100270). J.L. is supported by a Ph.D. scholarship from the China Scholarship Council and the Norma Hilda Schuster Scholarship; A.S. is supported by and Australian Postgraduate Award. M.L. was supported by the Brutton Bequest. Ethan Goddard is thanked for valuable discussions, and Alexandre Tolotchov is thanked for assistance with the construction and use of the MicrobeMeter. We thank the Melbourne Mass Spectrometry and Proteomics Facility of The University of Melbourne.

## Notes

https://www.ebi.ac.uk/pride/

### Competing Interest Statement

The authors have declared no competing interest.

### Summary of Updates

General revisions across entire manuscript. New data on a candidate SLA dehydrogenase.

## References

1. Durham BP. 2021. Deciphering metabolic currencies that support marine microbial networks. mSystems 6:e0076321.

2. Durham BP, Boysen AK, Carlson LT, Groussman RD, Heal KR, Cain KR, Morales RL, Coesel SN, Morris RM, Ingalls AE, Armbrust EV. 2019. Sulfonate-based networks between eukaryotic phytoplankton and heterotrophic bacteria in the surface ocean. Nat Microbiol 4:1706–1715.

3. Moran MA, Durham BP. 2019. Sulfur metabolites in the pelagic ocean. Nat Rev Microbiol 17:665–678.

4. Chaudhary S, Sindhu SS, Dhanker R, Kumari A. 2023. Microbes-mediated sulphur cycling in soil: Impact on soil fertility, crop production and environmental sustainability. Microbiol Res 271:127340.

5. Shaw DK, Sekar J, Ramalingam PV. 2022. Recent insights into oceanic dimethylsulfoniopropionate biosynthesis and catabolism. Environ Microbiol 24:2669–2700.

6. Thume K, Gebser B, Chen L, Meyer N, Kieber DJ, Pohnert G. 2018. The metabolite dimethylsulfoxonium propionate extends the marine organosulfur cycle. Nature 563:412–415.

7. Celik E, Maczka M, Bergen N, Brinkhoff T, Schulz S, Dickschat JS. 2017. Metabolism of 2,3-dihydroxypropane-1-sulfonate by marine bacteria. Org Biomol Chem 15:2919–2922.

8. Cook AM, Denger K, Smits TH. 2006. Dissimilation of C3-sulfonates. Arch Microbiol 185:83–90.

9. Goddard-Borger ED, Williams SJ. 2017. Sulfoquinovose in the biosphere: occurrence, metabolism and functions. Biochem J 474:827–849.

10. Benning C. 1998. Biosynthesis and function of the sulfolipid sulfoquinovosyl diacylglycerol. Annu Rev Plant Physiol Plant Mol Biol 49:53–75.

11. Snow AJD, Burchill L, Sharma M, Davies GJ, Williams SJ. 2021. Sulfoglycolysis: catabolic pathways for metabolism of sulfoquinovose. Chem Soc Rev 50:13628–13645

12. Wei Y, Tong Y, Zhang Y. 2022. New mechanisms for bacterial degradation of sulfoquinovose. Biosci Rep 42:BSR20220314.

13. Kaur A, Pickles IB, Sharma M, Madeido Soler N, Scott NE, Pidot SJ, Goddard-Borger ED, Davies GJ, Williams SJ. 2023. Widespread Family of NAD^+^-Dependent Sulfoquinovosidases at the Gateway to Sulfoquinovose Catabolism. J Am Chem Soc 145:28216–28223.

14. 14. Speciale G, Jin Y, Davies GJ, Williams SJ, Goddard-Borger ED. 2016. YihQ is a sulfoquinovosidase that cleaves sulfoquinovosyl diacylglyceride sulfolipids. Nat Chem Biol 12:215–217.

15. Abayakoon P, Jin Y, Lingford JP, Petricevic M, John A, Ryan E, Wai-Ying Mui J, Pires DEV, Ascher DB, Davies GJ, Goddard-Borger ED, Williams SJ. 2018. Structural and biochemical insights into the function and evolution of sulfoquinovosidases. ACS Cent Sci 4:1266–1273.

16. Frommeyer B, Fiedler AW, Oehler SR, Hanson BT, Loy A, Franchini P, Spiteller D, Schleheck D. 2020. Environmental and intestinal phylum Firmicutes bacteria metabolize the plant sugar sulfoquinovose via a 6-deoxy-6-sulfofructose transaldolase pathway. iScience 23:101510.

17. Denger K, Weiss M, Felux AK, Schneider A, Mayer C, Spiteller D, Huhn T, Cook AM, Schleheck D. 2014. Sulphoglycolysis in Escherichia coli K-12 closes a gap in the biogeochemical sulphur cycle. Nature 507:114–117.

18. Felux AK, Spiteller D, Klebensberger J, Schleheck D. 2015. Entner-Doudoroff pathway for sulfoquinovose degradation in Pseudomonas putida SQ1. Proc Natl Acad Sci USA 112:E4298–305.

19. Liu J, Wei Y, Ma K, An J, Liu X, Liu Y, Ang EL, Zhao H, Zhang Y. 2021. Mechanistically diverse pathways for sulfoquinovose degradation in bacteria. ACS Catalysis 11:14740–14750.

20. Liu Y, Wei Y, Zhou Y, Ang EL, Zhao H, Zhang Y. 2020. A transaldolase-dependent sulfoglycolysis pathway in Bacillus megaterium DSM 1804. Biochem Biophys Res Commun 533:1109–1114.

21. Weinitschke S, Hollemeyer K, Kusian B, Bowien B, Smits TH, Cook AM. 2010. Sulfoacetate is degraded via a novel pathway involving sulfoacetyl-CoA and sulfoacetaldehyde in Cupriavidus necator H16. J Biol Chem 285:35249–54.

22. Denger K, Huhn T, Hollemeyer K, Schleheck D, Cook AM. 2012. Sulfoquinovose degraded by pure cultures of bacteria with release of C_3_-organosulfonates: complete degradation in two-member communities. FEMS Microbiol Lett 328:39–45.

23. Tong Y, Wei Y, Hu Y, Ang EL, Zhao H, Zhang Y. 2019. A pathway for isethionate dissimilation in Bacillus krulwichiae. Appl Environ Microbiol 85.

24. Ruff J, Denger K, Cook AM. 2003. Sulphoacetaldehyde acetyltransferase yields acetyl phosphate: purification from Alcaligenes defragrans and gene clusters in taurine degradation. Biochem J 369:275–85.

25. Denger K, Cook AM. 2010. Racemase activity effected by two dehydrogenases in sulfolactate degradation by Chromohalobacter salexigens: purification of (S)-sulfolactate dehydrogenase. Microbiology 156:967–974.

26. Wei Y, Zhang Y. 2021. Glycyl radical enzymes and sulfonate metabolism in the microbiome. Annu Rev Biochem 90:817–846.

27. Sharma M, Lingford JP, Petricevic M, Snow AJP, Zhang Y, Jarva, M., , Mui JW-Y, Scott NE, Saunders EC, Mao R, Epa R, da Silva BM, Pires DEV, Ascher DB, McConville MJ, Davies, G.J., , Williams SJ, Goddard-Borger, E.D. 2022. Oxidative desulfurization pathway for complete catabolism of sulfoquinovose by bacteria. Proc Natl Acad Sci USA 119:e2116022119.

28. Liem PQ, Laur M-H. 1976. Structures, teneurs et compositions des esters sulfuriques, sulfoniques, phosphoriques des glycosyldiglycérides de trois fucacées. Biochimie 58:1367–1380.

29. Vinogradov E, Deschatelets L, Lamoureux M, Patel GB, Tremblay TL, Robotham A, Goneau MF, Cummings-Lorbetskie C, Watson DC, Brisson JR, Kelly JF, Gilbert M. 2012. Cell surface glycoproteins from Thermoplasma acidophilum are modified with an N-linked glycan containing 6-C-sulfofucose. Glycobiology 22:1256–67.

30. Jain C, Rodriguez-R LM, Phillippy AM, Konstantinidis KT, Aluru S. 2018. High throughput ANI analysis of 90K prokaryotic genomes reveals clear species boundaries. Nat Commun 9:5114.

31. Rodriguez RLM, Conrad Roth E, Viver T, Feistel Dorian J, Lindner Blake G, Venter Stephanus N, Orellana Luis H, Amann R, Rossello-Mora R, Konstantinidis Konstantinos T. 2023. An ANI gap within bacterial species that advances the definitions of intra-species units. mBio 15:e02696–23.

32. Figueras María J, Beaz-Hidalgo R, Hossain Mohammad J, Liles Mark R. 2014. Taxonomic affiliation of new genomes should be verified using average nucleotide identity and multilocus phylogenetic analysis. Genome Announcements 2:10.1128/genomea.00927-14.

33. Mui JW-Y, Souza DPD, Saunders EC, McConville MJ, Williams SJ. 2023. Remodeling of Carbon Metabolism during Sulfoglycolysis in Escherichia coli. Appl Environ Microbiol 89:e02016–22.

34. Weinitschke S, Denger K, Cook AM, Smits THM. 2007. The DUF81 protein TauE in Cupriavidus necator H16, a sulfite exporter in the metabolism of C2 sulfonates. 153:3055-3060.

35. Rein U, Gueta R, Denger K, Ruff J, Hollemeyer K, Cook AM. 2005. Dissimilation of cysteate via 3-sulfolactate sulfo-lyase and a sulfate exporter in Paracoccus pantotrophus NKNCYSA. Microbiology 151:737–747.

36. Burrichter A, Denger K, Franchini P, Huhn T, Müller N, Spiteller D, Schleheck D. 2018. Anaerobic Degradation of the Plant Sugar Sulfoquinovose Concomitant With H2S Production: Escherichia coli K-12 and Desulfovibrio sp. Strain DF1 as Co-culture Model. Front Microbiol 9:2792.

37. Burchill L, Sharma M, Soler NM, Goddard-Borger ED, Davies GJ, Williams SJ. 2025. Structure, kinetics, and mechanism of Pseudomonas putida sulfoquinovose dehydrogenase, the first enzyme in the sulfoglycolytic Entner-Doudoroff pathway. Biochem J 482:57–72.

38. Zallot R, Oberg N, Gerlt JA. 2019. The EFI web resource for genomic enzymology tools: leveraging protein, genome, and metagenome databases to discover novel enzymes and metabolic pathways. Biochemistry 58:4169–4182.

39. Oberg N, Zallot R, Gerlt JA. 2023. EFI-EST, EFI-GNT, and EFI-CGFP: Enzyme Function Initiative (EFI) Web Resource for Genomic Enzymology Tools. J Mol Biol 435:168018.

40. Mikosch CARM, Denger K, Schäfer E-M, Cook AM. 1999. Anaerobic oxidations of cysteate: degradation via L-cysteate: 2-oxoglutarate aminotransferase in Paracoccus pantotrophus. Microbiology 145:1153–1160.

41. Li J, Epa R, Scott NE, Skoneczny D, Sharma M, Snow AJD, Lingford JP, Goddard-Borger ED, Davies GJ, McConville MJ, Williams SJ. 2020. A Sulfoglycolytic Entner-Doudoroff Pathway in Rhizobium leguminosarum bv. trifolii SRDI565. Appl Environ Microbiol 86:e00750–20.

42. Snow AJD, Sharma M, Lingford JP, Zhang Y, Mui JW, Epa R, Goddard-Borger ED, Williams SJ, Davies GJ. 2022. The sulfoquinovosyl glycerol binding protein SmoF binds and accommodates plant sulfolipids. Curr Res Struct Biol 4:51–58.

43. Arumapperuma T, Snow AJD, Lee M, Sharma M, Zhang Y, Lingford JP, Goddard-Borger ED, Davies GJ, Williams SJ. 2024. Capture-and-release of a sulfoquinovose-binding protein on sulfoquinovose-modified agarose. Org Biomol Chem 22:3237–3244.

44. Sharma M, Abayakoon P, Epa R, Jin Y, Lingford JP, Shimada T, Nakano M, Mui JWY, Ishihama A, Goddard-Borger ED, Davies GJ, Williams SJ. 2021. Molecular basis of sulfosugar selectivity in sulfoglycolysis. ACS Cent Sci 7:476–487.

45. De Ley J, Doudoroff M. 1957. The metabolism of D-galactose in Pseudomonas saccharophila. J Biol Chem 227:745–57.

46. Yun EJ, Lee S-H, Kim S, Ryu HS, Kim KH. 2024. Catabolism of 2-keto-3-deoxy-galactonate and the production of its enantiomers. Appl Microbiol Biotechnol 108:403.

47. Li J, Sharma M, Meek R, Alhifthi A, Armstrong Z, Soler NM, Lee M, Goddard-Borger ED, Blaza JN, Davies GJ, Williams SJ. 2023. Molecular basis of sulfolactate synthesis by sulfolactaldehyde dehydrogenase from Rhizobium leguminosarum. Chem Sci 14:11429–11440.

48. Roy AB, Hewlins MJ, Ellis AJ, Harwood JL, White GF. 2003. Glycolytic breakdown of sulfoquinovose in bacteria: a missing link in the sulfur cycle. Appl Environ Microbiol 69:6434–6441.

49. Abayakoon P, Epa R, Petricevic M, Bengt C, Mui JWY, van der Peet PL, Zhang Y, Lingford JP, White JM, Goddard-Borger ED, Williams SJ. 2019. Comprehensive synthesis of substrates, intermediates and products of the sulfoglycolytic Embden-Meyerhoff-Parnas pathway. J Org Chem 84:2901–2910.

50. Graham DE, Xu H, White RH. 2002. Identification of coenzyme M biosynthetic phosphosulfolactate synthase: a new family of sulfonate-biosynthesizing enzymes. J Biol Chem 277:13421–13429.

51. Sasidharan K, Martinez-Vernon AS, Chen J, Fu T, Soyer OS. 2018. A low-cost DIY device for high resolution, continuous measurement of microbial growth dynamics. bioRxiv doi:10.1101/407742:407742.

52. Chaumeil PA, Mussig AJ, Hugenholtz P, Parks DH. 2019. GTDB-Tk: a toolkit to classify genomes with the Genome Taxonomy Database. Bioinformatics (Oxford, England) 36:1925–7.

53. Parks DH, Chuvochina M, Chaumeil PA, Rinke C, Mussig AJ, Hugenholtz P. 2020. A complete domain-to-species taxonomy for Bacteria and Archaea. Nat Biotechnol 38:1079–1086.

54. Yoon SH, Ha SM, Lim J, Kwon S, Chun J. 2017. A large-scale evaluation of algorithms to calculate average nucleotide identity. Antonie Van Leeuwenhoek 110:1281–1286.

55. Blakeney AB, Mutton LL. 1980. A simple colorimetric method for the determination of sugars in fruit and vegetables. J Sci Food Agric 31:889–897.

56. Sörbo B. 1987. Sulfate: Turbidimetric and nephelometric methods, p 3-6, Methods Enzymol, vol 143. Academic Press.

57. Brychkova G, Yarmolinsky D, Fluhr R, Sagi M. 2012. The determination of sulfite levels and its oxidation in plant leaves. Plant Sci 190:123–130.

58. Kurmanbayeva A, Brychkova G, Bekturova A, Khozin I, Standing D, Yarmolinsky D, Sagi M. 2017. Determination of Total Sulfur, Sulfate, Sulfite, Thiosulfate, and Sulfolipids in Plants, p 253–271. In Sunkar R (ed), Plant Stress Tolerance: Methods and Protocols doi:10.1007/978-1-4939-7136-7_15. Springer New York, New York, NY.

59. Kulak NA, Pichler G, Paron I, Nagaraj N, Mann M. 2014. Minimal, encapsulated proteomic-sample processing applied to copy-number estimation in eukaryotic cells. Nat Methods 11:319–24.

60. Harney DJ, Hutchison AT, Hatchwell L, Humphrey SJ, James DE, Hocking S, Heilbronn LK, Larance M. 2019. Proteomic Analysis of Human Plasma during Intermittent Fasting. J Proteome Res 18:2228–2240.

61. Rappsilber J, Mann M, Ishihama Y. 2007. Protocol for micro-purification, enrichment, pre-fractionation and storage of peptides for proteomics using StageTips. Nat Protoc 2:1896–1906.

62. Cox J, Mann M. 2008. MaxQuant enables high peptide identification rates, individualized p.p.b.-range mass accuracies and proteome-wide protein quantification. Nat Biotechnol 26:1367–1372.

63. Tyanova S, Temu T, Sinitcyn P, Carlson A, Hein MY, Geiger T, Mann M, Cox J. 2016. The Perseus computational platform for comprehensive analysis of (prote)omics data. Nat Methods 13:731.

64. Shannon P, Markiel A, Ozier O, Baliga NS, Wang JT, Ramage D, Amin N, Schwikowski B, Ideker T. 2003. Cytoscape: A Software Environment for Integrated Models of Biomolecular Interaction Networks. Genome Res 13:2498–2504.

65. Perez-Riverol Y, Csordas A, Bai J, Bernal-Llinares M, Hewapathirana S, Kundu DJ, Inuganti A, Griss J, Mayer G, Eisenacher M, Pérez E, Uszkoreit J, Pfeuffer J, Sachsenberg T, Yilmaz S, Tiwary S, Cox J, Audain E, Walzer M, Jarnuczak AF, Ternent T, Brazma A, Vizcaíno JA. 2019. The PRIDE database and related tools and resources in 2019: improving support for quantification data. Nucleic Acids Res 47:D442–d450.

