## Supplementary Figures for "Tandem sulfofucolytic-sulfolactate sulfolyase pathway for catabolism of the rare sulfosugar sulfofucose"

SUPPLEMENTAL FIGURES

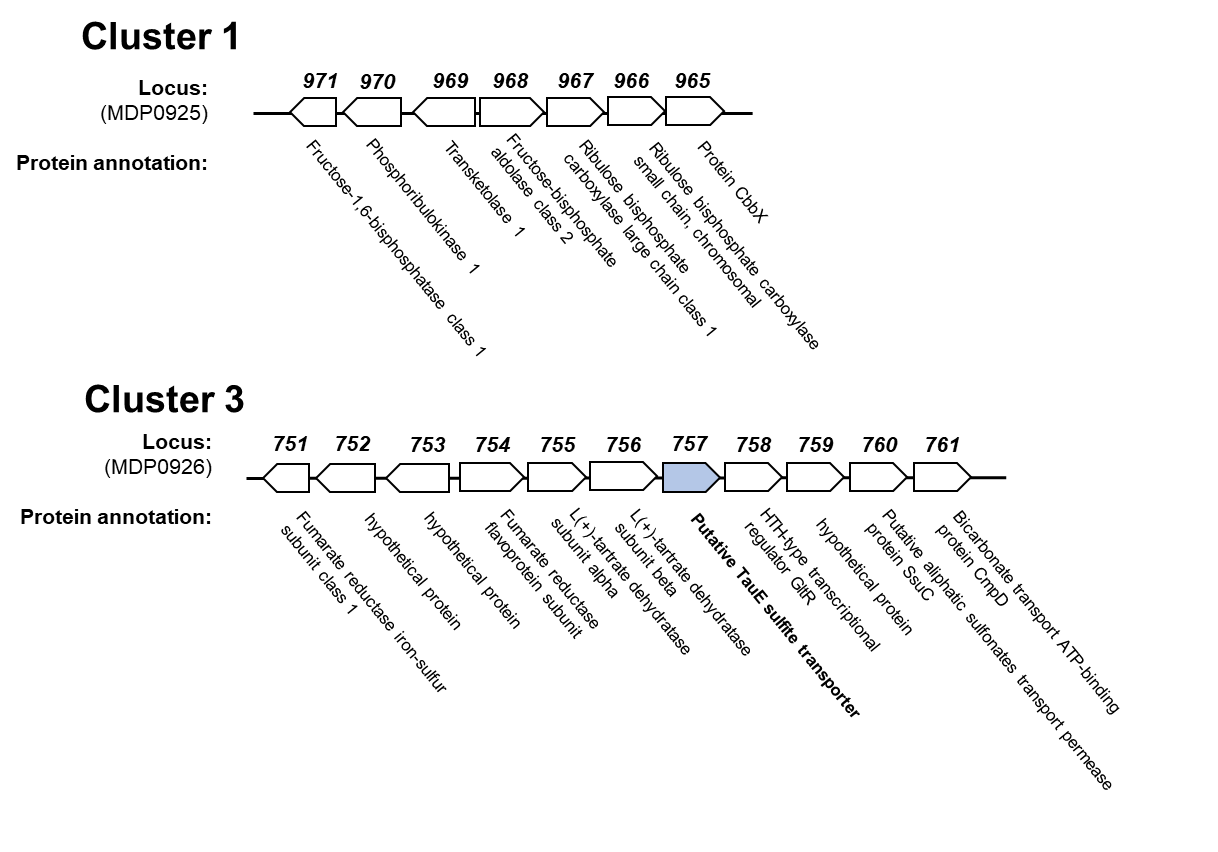

**Figure S1.** Gene clusters 1 and 3 identified by comparative proteomics. Annotations were added automatically during assembly and annotation, except for annotation for TauE added manually.
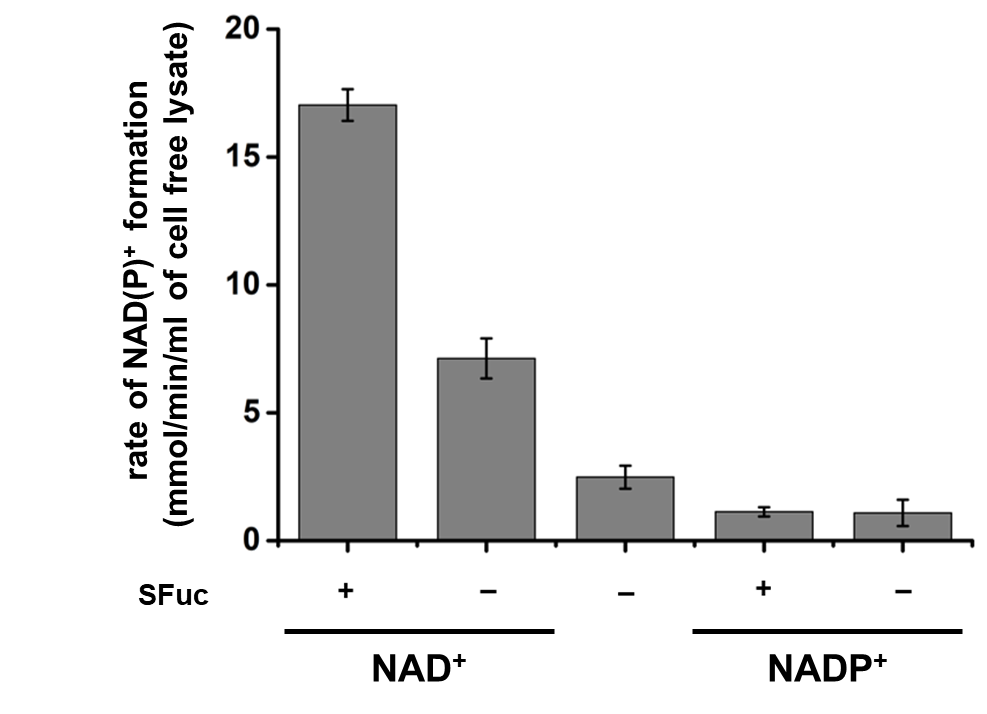

**Figure S2.** NAD^+^-dependent sulfofucose dehydrogenase activity of cell free lysate of *P. wurundjeri* strain Merri. Reactions contained 1.1 mM SFuc, 6.5 mM MgCl_2_, and 0.5 mM NAD^+^ or NADP^+^ in Tris-HCl (100 mM, pH 7.5). Reactions were initiated by the addition of 20 μL of strain Merri cell lysate and absorbance was measured at 340 nm.

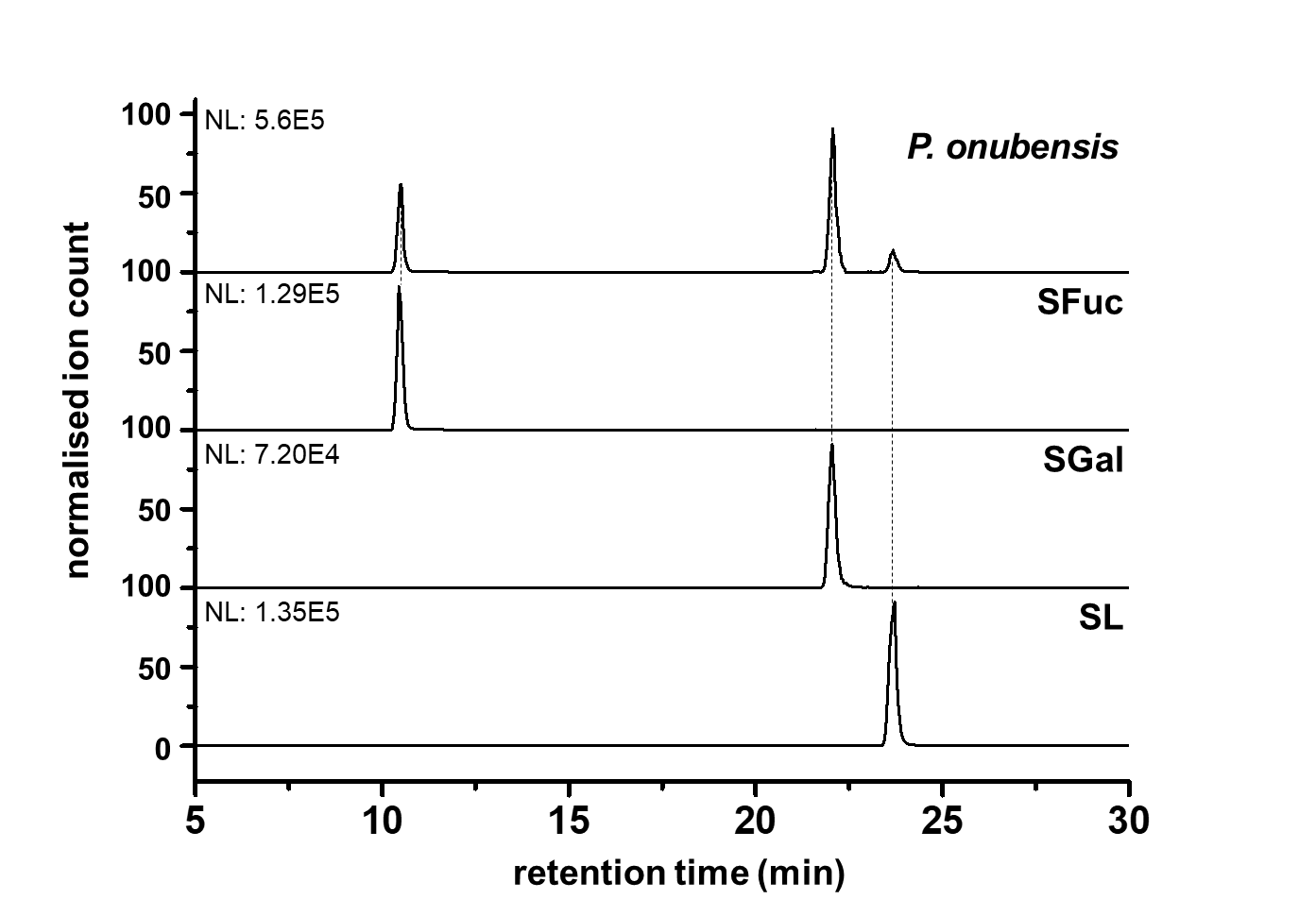

**Figure S3. Ion chromatography-mass spectrometry analysis of *P. wurundjeri*** **strain Merri grown on SFuc shows SGal and SL as metabolites.** Extracted ion chromatographs of metabolites from *P. wurundjeri* (top), and chemically synthesised SFuc, SGal and SL.

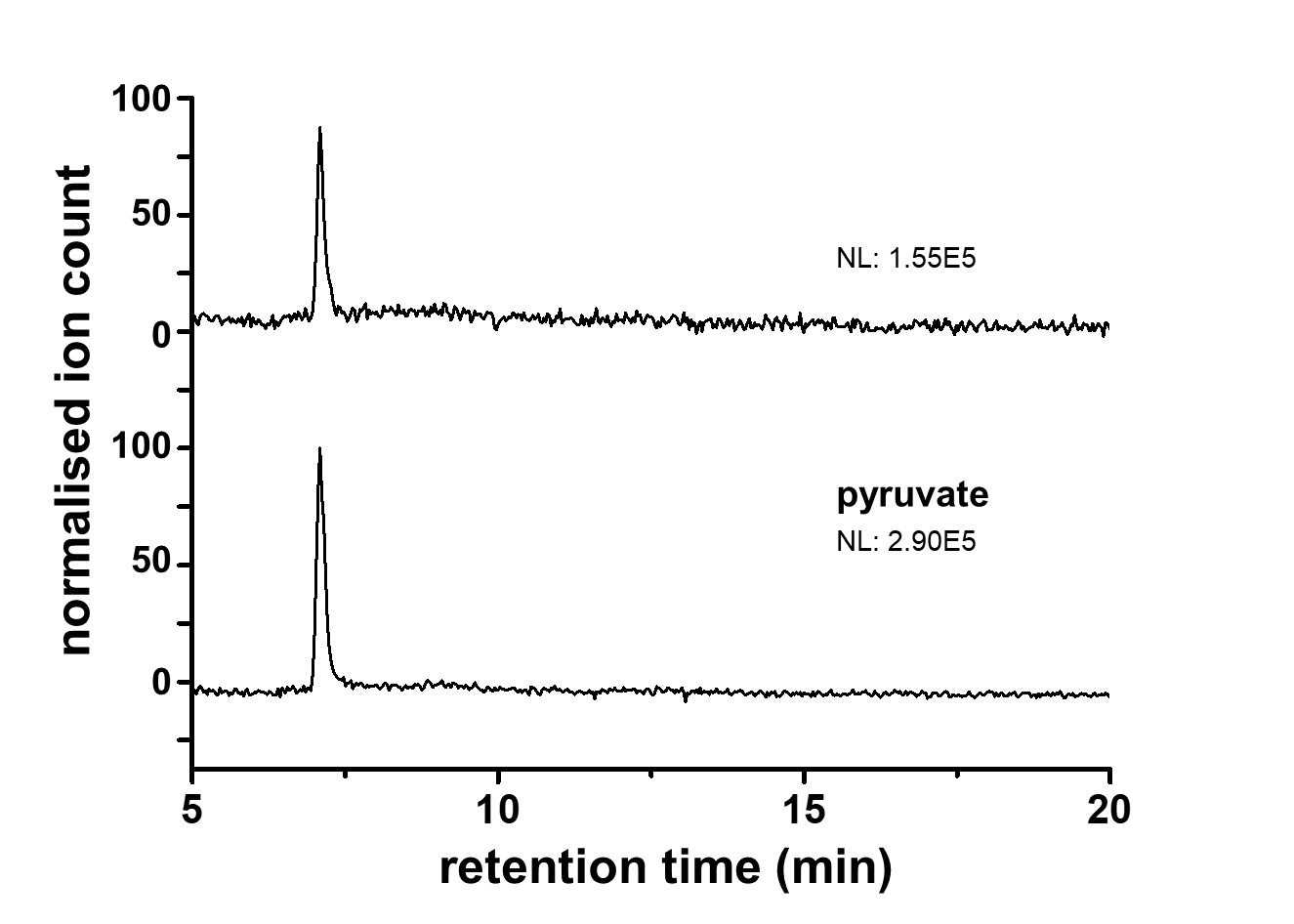

**Figure S4.** **Pyruvate is generated by KDSGal aldolase (SfcE).** Targeted SIM IC-MS extracted ion chromatographs monitoring for *m/z* 87 (pyruvate) showing (top) reaction mixture containing KDSGal, MgCl_2_ and SfcE and (bottom) a pyruvate standard.

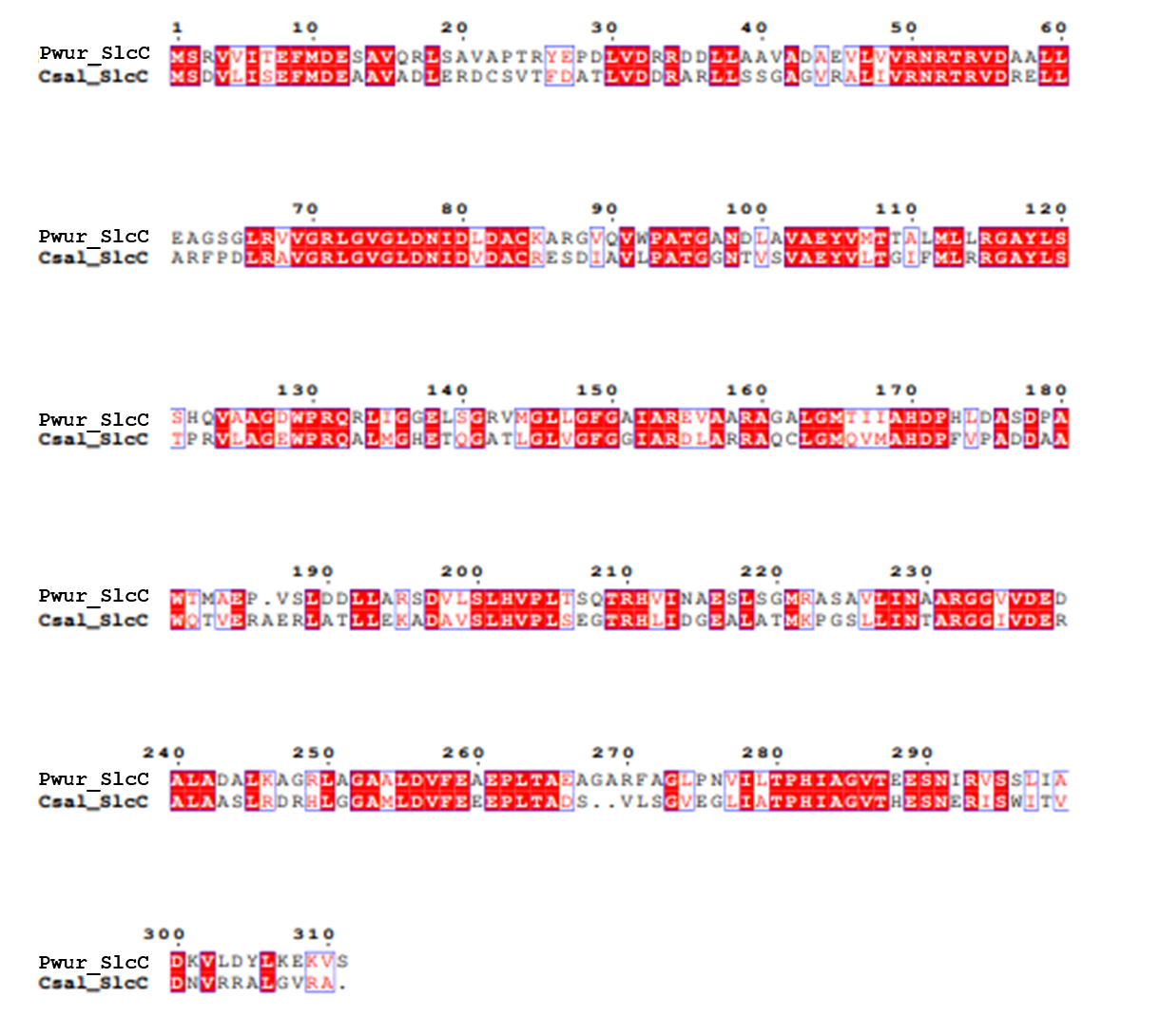

**Figure S5.** Sequence alignment of proposed SlcC protein from *Paracoccus wurundjeri* strain Merri and biochemically validated SlcC from *Chromohalobacter salexigens*.

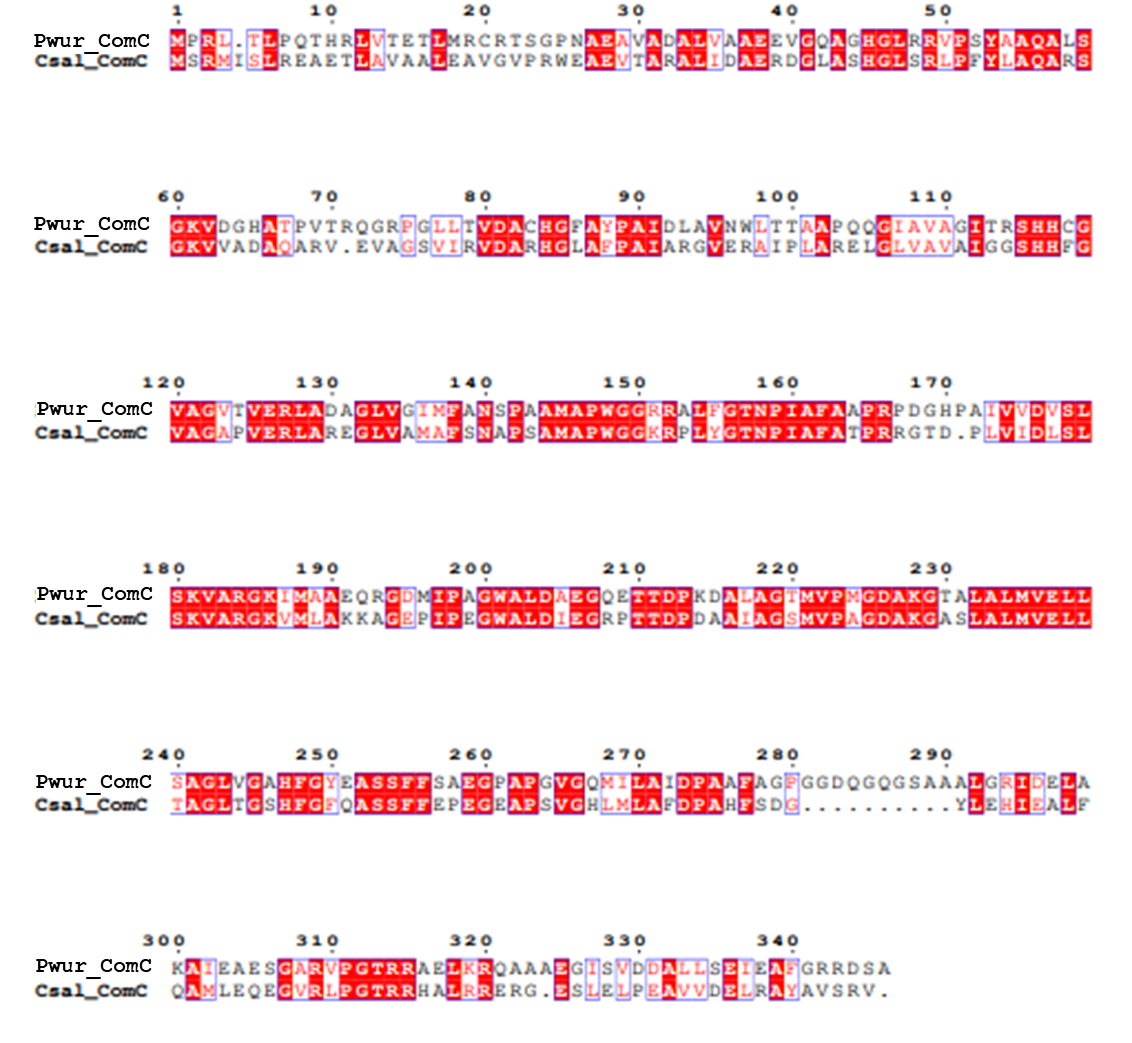

**Figure S6.** Sequence alignment of proposed ComC protein from *Paracoccus wurundjeri* strain Merri and biochemically validated ComC from *Chromohalobacter salexigens*.

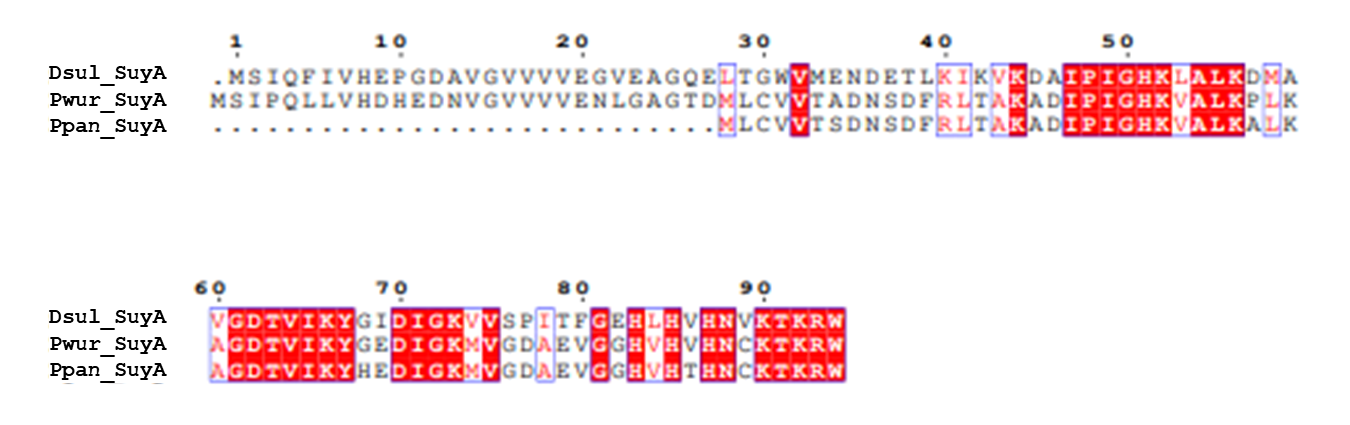

**Figure S7.** Sequence alignment of proposed SuyA protein from *Paracoccus wurundjeri* strain Merri, and biochemically validated SuyA from *Paracoccus pantotrophus* NKNCYSA and *Desulfovibrio sp. strain* DF1.

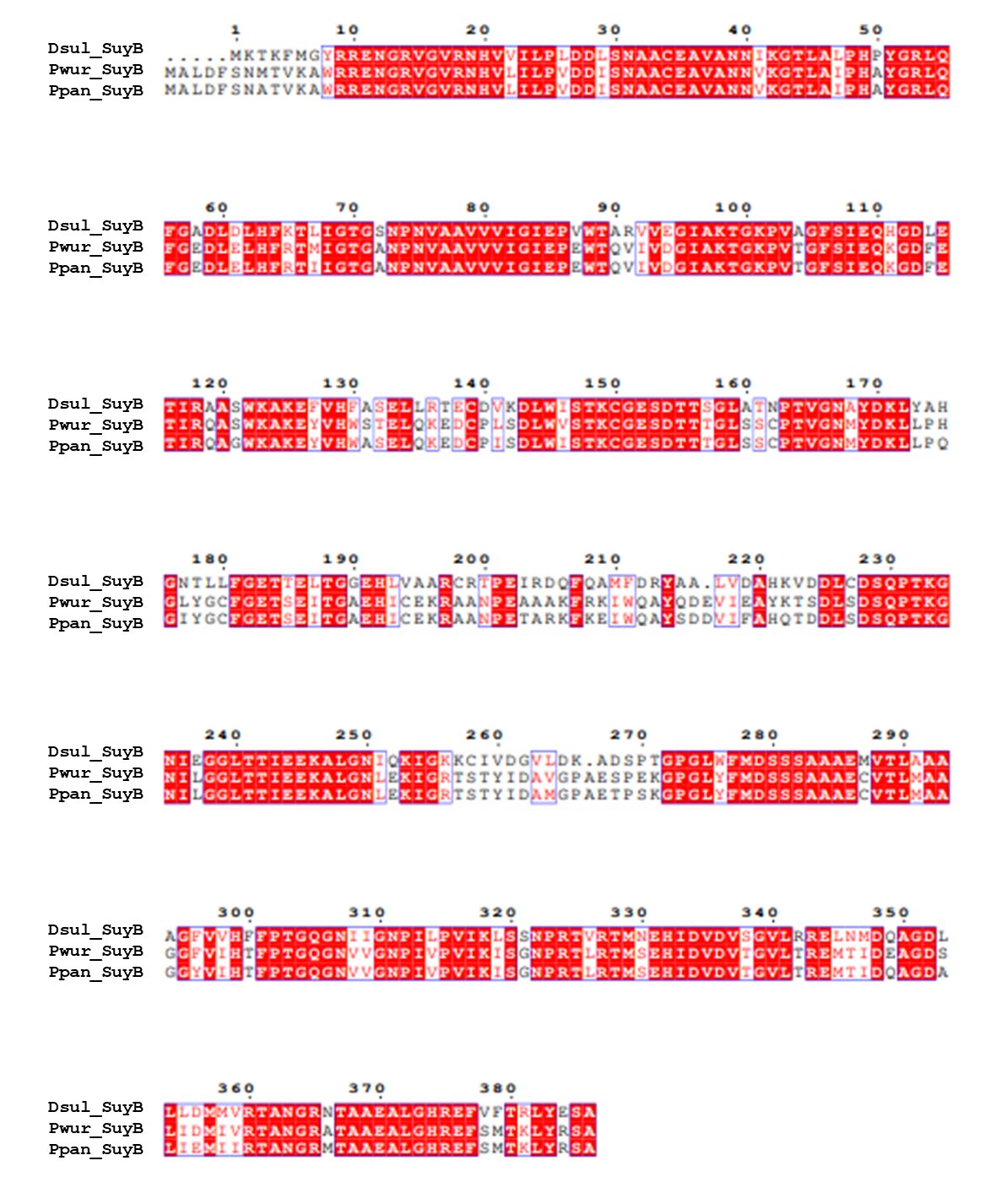

**Figure S8.** Sequence alignment of proposed SuyB protein from *Paracoccus wurundjeri* strain Merri, and biochemically validated SuyA from *Paracoccus pantotrophus* NKNCYSA and *Desulfovibrio sp. strain* DF1.
