## Supplementary Gene and Protein Sequences for "Tandem sulfofucolytic-sulfolactate sulfolyase pathway for catabolism of the rare sulfosugar sulfofucose"

SUPPLEMENTAL INFORMATION – GENE SEQUENCES

**Codon harmonised amino acid sequences of expressed *P. wurundjeri* SfcD, SfcE, SfcF, SfcH, and MDP0926266**

> SfcD (lactonase)_codon harmonised

ATGCCTCGGGTCATTGACATTTGGCAAGTGGAGCGCCCTGAGCTCCCTCGTTCGCTCCTGGGTGAGACGGTGCGCTGGCACGCTCGATCAGGCCGTGTTATATACGCTGACATAATGGGCCGCAAGTTACTGGCATACGACCCATTGACTGGACAACAAGACGACTGGAGTTTTGAAGCACCAGTTGGTGGTTGGTTCCCTACTATTCGTGACGAGATAATTCTGGCTGTGGGCCAAAATCTTGATTGGTTTGATATGAAGACCGGGTCACGTCGTCCACTGGCCAGTCTCGTTGGTGAAGACGCAGACATACGATTTAACGATGGCAGATGCGATCCCGCAGGTCGCTTGTGGATAAGCACCATGTCCATGTCTGGCAACCCACCGCCGCCTCGTGGAAGACTTTTCCGCTTGGATCCACCAGGTATTCTTACCCCTGTAATTGATGGATTACGTATACCAAACACATTAGCGTGGTCACCTGACGGCAATAGTATGTACTTTGCTGACTCTCTTACTCGGGAGACAGGGCGTTATGGTTACGACCCTCAGAATGGGGAACTGGGAACGCGCTCTGTGCTCTTTAGATTCAGTGACGCCGTCAGTGGTATACCAGACGGCGCATGTGTAGACGAAGAGGGTGGTATATGGATCGCAGTCCCACGTGGGTCTCGTGTAGAGCGTCGTATGCCAGATGGAAGCCTTGATACCGTCATACTTCTTCCTGCAGAGCGTCCCACGATGTGCGCATTCGGTGGGGCAGATCGAGACATTTTGTTTATTACATCCCAAAGCTTGTTTCTTAGCGAGGAAGAAAGACGCTTTCGCGTAAACGATGGGGCTTTCTTTGCGGTTAGACCGGGTGTTACTGGGCTCCCCGAGGCACAGTTTGATCCTGATACAATTCGGAATTAA

> SfcE (aldolase)_codon harmonised

ATGACTACTAGAAATGACTTTGCTAACCCCTTTCGTCGCGCACTCGCTGGCAAGAAAGTTTTAACGGGAATATGGAGTATGTTAAACTCTACGAACGCCATTGAAGGACTGGGATGGGCTGGGTTTGATTGGCTCGTTATTGACGCGGAGCACTCACCTGTGTCATTGCACGATGCTATGGCCCACTTACGGGCTTTGGGTGCAACCCCCACCATACCCGTTGTCCGCCTTCCGTGGAATGACAATGTACTCATTAAACACTACTTGGACATAGGAGCTCAAACAATAATGTTGCCCCTTATTCAAAACGCGCGTGAGGCTCAAGCAGCAGTACGTGCAATGCGGTATCCCCCGGGTGGTATGCGGGGCTTTGCAGCAATGCACAGAGCATCACGATATGGTCACATACCACAGTACGTCGAGAGAGCGGAAGAGACTTTATTTATGATTGTCCAAGTCGAAACTGTAGAAGCCCTCGACCGCTTAGATGAAATTGCCAGCGTTGAGGGAGTTGATGCTGTCTTCTTCGGTCCGGGAGACTTAAGTGCGTCGATGGGCCTGTTAGGGAAACCAGGCCAAACTGAGGTATTTGATCGTATTGTGGAAGCCTCAAAGACAGTTCAACGACTGGGAAAATCGACTGGAGTGTTGGCACCGTCCATTGGGCATGCAAAGTCTTACTGTTCGGCAGGAGTCAACTTTGTTTCTGTCGCTACGGACTGTGCATTGCTCTTCCGCAATGCGGATGCCTTAGCAGCAGACTTTGCTGCTTTTACAGCAGCAGCCACCTGA

> SfcF (dehydratase)_codon harmonised

ATGGATCGCATTGCAGAGATTACTAGTTTTACAGTACCACCACGTTGGATCTTTGTGCGCGTTCGGACGGCAGAGGGATTGACAGGCTGGGGTGAAGCGATTATTCCTAAACGTCGGAACGCAGTGATAGGGGCAATACGAGACTTGACTCAAGTGGTTTTAGGAATGGACCCGGCTCGTATTGAAGACATTGCCAGTAGCCTGAGAAAAGGTTCATTCTTTCGCAATGGACCTATTCTCGGCACTGCGATCGCTGCCGTGGAGATCGCTCTGTGGGACATTAAAGGCCAAAGAGCTGGCCTGCCTGTGTTTGAATTCCTTGGCGGTCGTGTCAGAGACAACATAAGAAGCTACACGTGGATAGGTGGGGATTCTCCAGCTAACGTTGTATCTCATGCCAAGGAGAGAGTTGAGCAAGGATTTGACGCTGTGAAAATGAATGCCACACCGGCGGTTGCACACTTAGAGTGGCGTGAAGCCACTGAGAATTTAGTACAGCGAATGGGGTCTCTTCGTGACGCTTTTGGTGGTAGCATAGATATCGCCCTTGACTTTCACGGTCGTGTGCCTAGAAGTGTTCTGAAGCAAATGGTAAAAGAGATAGAACCATTTGACCCATTGTGGATTGAGGAACCCTTTCTTCCGGAGCACGTTGGTGCTGACGAATTAATGGCCCGAATTTGTCCACACATACCAATAGCGACGGGAGAGAGATTGCTTCACCGCTGGGACTTTCAAAGACTGTTAGAGAGAGGTGGGGTTGACGTTATACAACCGGATATCTCGATAACCGGTCTGTTTGAAATGGAGAAAATTGCAAGACTTGCAGAAATCTATGATGTGGGTGTGGCACCTCACTGCCCTAACGGGCCAATAAGCTTGGCGGCAAGTCTCCAAGTCGACTTTTGTTGTGCGAACACCGTTATCCAAGAACAGTCTCTGGGCTTGCATTATAACCAAGGGTACGCAGGGTTGCCACCAGCTGATATTCTTGACTACATAGGCGACCCGGACGTTCTCACAACTCGCAATGGGCGTTTTGATTGCCCGAGTGCACCAGGGCTTGGTTTAATATTGAAGGGTGACACCATTGAAGCTGCCCACACTGATTGGAGTTTACCGGACCCTGATTGGCGACACAGAGATGGTGTATACGCTGAGTGGTAA

> SfcH (dehydrogenase)_codon harmonised

ATGACAGATATCATAACTGAGCCCGGCGTTCTTGTCACCGGTGCTGCCGGTGGTATAGGTCGTGCTGTTGCAGGTTTATTTGCTGATCAGGGGCATGCTGTAACACTGACGGACAGAGACTCGGCAGGTCTTGAGGATATAGGCGGACGGTTGCGTGATCGCGGGTGCAAGGTAGATATGATTGCTCAGGATCTTGCCGATCCCGACGCACCTGCCATGTTAGTCGCCTGCACCGTCGAGAAATGGGGCGGTATTGGAGTTCTTGTGAACAATGCTGCGCATCATGGTAAACGACAGAGTGTGCTGTCGAGCGAACCAGACGAATGGCGCCAGGTTTTCGAGGTGAATGTGATAGCCGCAGCAGCGCTGGCACGGTATGCAGCTCTCGACATGGCACGTCGCAAAACGGGTGCGATCATCAATGTGGGGTCCATACAACAGGCATTGCCGGTTAGTTCCTATGCCGCATATGTCGCAAGCAAGGGTGCAGTAGCCGGGATGACTCGGGCACTTGCCGTGGAATTGGGTCTGCTGGGCATAAGAGTCAACAGTGTCAGCCCTGGTGTGATCGGCACGGAGAATTTCCGCCGTGAATTGGAAACCCGTCACGATGGCGCTTCAGCTATCAGTTATCCGTCTTTATTGGGACGCCTTGGAACGCCTGAGGATGTGGCTCATGTAATCGCGTTTCTCGCCGGTCCCCATTCTTCTCATGTAACTGGTGCAGATTACGTGGTTGATGGCGGACGTGGGATCTCACGTCGCACCGACCCTTTCCATGCCGAAATCACACATCCAGTGAGTGGTAAATAG

> MDP0926266 (Putative threonine dehydrogease tested for SLADH activity)_codon harmonised

ATGATCCCTAATGAGAAAACCATGCAAGCTGCAGACTTCTTGGGAGAGGATCGTATTGAAATAGTATCACGACCACTTCCAGAGCCTGCAGAGGGCGAGGTGTTGCTCAGAGTGGCCGCCAACGCGCTCTGTGGTTCTGACTTAAAGCTCTGGCACGCTGGGGCCCAACATATAGCTGGTCATGAAATTGCTGGTTGGGTCCAACAGCCAGGCCACCCCTTGAATGGACAACTCTGTGCAGTATACATACCCTTACACTGTGGGGAGTGTGCTGTATGCCTCAGAGGGGACACACAGTCTTGTATAACAGTTTCAAGCTTAATTGGTTGGAATAGAGACGGTGGGTACGCACAGTACTTGACGGTTCCCGAGAATTGTTTACTCCCCGTACCCGGAGACATTGACGCTGCATTAGCTCCTTTGCTCTTAGACACAATAGGGACCTCAGCACACGCACTTCGAGAAGCGTCTCGTCACCTGGCTACTGATGCACCAAGTGTGTTAGTAACTGGTGCTGGCCCCGTAGGCTTGGGTGTAGTACTTGCGGCGGCCGCCTTAGGTCACGCTCAAGTTGACGTTGCTGAACCCAACCCGGCACGGGCGGCTATAGCTCGCGAGTTTGGGGCAAACATTGTTCCTGTAGGATCACACGATAGAAGATACGACCTCATAGTTGAGTGCTCTGGTAACCACGCTGCTCGGGACCTTGCGATACACTTGGTGTTGCCGAAGGGTGTAATTGTCCTGGTTGGGGAGAATGCAGCCCCCTGGTCAGTGACAGAGGATAAAGTTTTCCGACGTAAAGACTTTGCATTACTCAGAACGTTTTACTTTCCACGTGACGACTTTGCGGCAAACGTTGAGCTTTTAAGAGCTAATCGTGAAAAGTACGCAAGACTTGTTGACGATGCTTTCCCTATTGCTGAGTTGCCTCAAAAGTTTGCAGACTTTGCTGAGGGTAAGAGTATTAAGCCGATATTAAGCTTTATTGGCGAACAGTAA
