## Supplementary Tables for "Tandem sulfofucolytic-sulfolactate sulfolyase pathway for catabolism of the rare sulfosugar sulfofucose"

SUPPLEMENTAL TABLES

**Table S1.** Protein family membership and proposed functions of proteins encoded by genes within cluster 2.

| **Locus**  (MDP0926) | **InterPro classification** | **PFAM** | **Proposed function** |
| --- | --- | --- | --- |
| 256 | IPR050679 | PF00392  PF07702 | Transcriptional regulator |
| 257 | IPR044144 | - | Sulfolactate lyase subunit alpha (SuyA) |
| 258 | IPR052172 | PF20629  PF04295 | Sulfolactate lyase subunit beta (SuyB) |
| 259 | IPR050857 | PF00389  PF02826 | *S*-sulfolactate dehydrogenase (SlcC) |
| 260 | IPR003767 | PF02615 | *R*-sulfolactate dehydrogenase (ComC) |
| 261 | None predicted | PF00356  PF13377 | Transcriptional regulator |
| 262 | IPR050490  IPR006059 | PF01547 |  |
| 263 | IPR002347 | PF13561 |  |
| 264 | IPR052730 | PF00528 |  |
| 265 | IPR050901 | PF00528 |  |
| 266 | IPR050129 | PF08240  PF00107 | Putative threonine dehydrogenase |
| 267 | IPR047641 | PF00005  PF08402 |  |
| 268 | None predicted | PF00294 |  |
| 269 | IPR050552)  IPR002915 | PF01791 |  |
| 270 | IPR046945 | PF13378  PF02746 |  |
| 271 | None predicted | PF07729  PF00392 | Transcriptional regulator |
| 272 | IPR006059 | PF01547 | putative sulfofucose binding protein (SfcK) |
| 273 | None predicted | PF00528 | putative ABC transporter protein (SfcJ) |
| 274 | IPR050901 | PF00528 | putative ABC transporter protein (SfcI) |
| 275 | IPR002347 | PF13561 | sulfofuconate dehydrogenase (SfcH) |
| 276 | IPR047641 | PF00005  PF08402 | putative ABC transporter protein (SfcG) |
| 277 | IPR034593 | PF13378  PF02746 | sulfogalactonate dehydratase (SfcF) |
| 278 | IPR050251 | PF03328 | 2-keto-3-deoxysulfogalactonate aldolase (SfcE) |
| 279 | IPR005511 | PF08450 | sulfofuconolactone lactonase (SfcD) |

**Table S2.** Protein family membership of enzymes involved in SL biomineralization from *C. salexigens*.^1^

| **Enzyme** | **PFAM** |
| --- | --- |
| *S*-sulfolactate dehydrogenase  SlcC | PF00389  PF02826 |
| *R*-sulfolactate dehydrogenase ComC | PF02615 |
| Sulfolactate lyase subunit alpha SuyA | - |
| Sulfolactate lyase subunit beta SuyB | PF20629  PF04295 |

**Table S3.** Protein family membership of sulfo-EMP enzymes from *E. coli* K12.^2^

| **Enzyme** | **PFAM** |
| --- | --- |
| Sulfoquinovose isomerase | PF07221 |
| Sulfofructose kinase | PF00294 |
| Sulfofructose-6-phosphate aldolase | PF01791 |

**Table S4.** Protein family membership of sulfo-ED enzymes from *P. putida* SQ1*.*^3^

| **Enzyme** | **PFAM** |
| --- | --- |
| Sulfoquinovose dehydrogenase SedA | PF13561 |
| Sulfogluconolactone lactonase SedB | PF08450 |
| Sulfogluconate dehydratase  SedC | PF00920 |
| 2-keto-3,6-dideoxy-6-sulfogluconate aldolase  SedD | PF03328 |
| Sulfolactaldehyde dehydrogenase GabD/SlaB | PF00171 |

**Table S5.** Uniprot sequence accession codes of Sfc*X* homologues of *P. wurundjeri* strain Merri sulfofucolytic proteins.

|  | **Taxon ID** | **SfcD** | **SfcE** | **SfcF** | **SfcG** | **SfcH** | **SfcI** | **SfcJ** | **SfcK** |
| --- | --- | --- | --- | --- | --- | --- | --- | --- | --- |
| ***Paracoccus alkanivorans* strain 4-2** | 2116655 | – | A0A3M0MEP9 | A0A3M0MGN3  A0A3M0MKR2 | A0A3M0MIZ6 | A0A3M0MFQ9 | A0A3M0MXY8 | A0A3M0MEX2 | A0A3M0MEQ1 |
| ***Rhizobiales* bacterium strain 65-79** | 1895815 | – | A0A1M2Z205 | A0A1M2Z1S0 | A0A1M2Z1Z4 | A0A1M2Z1W4 | A0A1M2Z1N3 | A0A1M2Z1U8 | A0A1M2Z1S6 |
| ***Rhizobiaceae* bacterium strain FW021_bin.52** | 1913961 | – | – | A0A522R0Q2 | A0A522R0Q6 | A0A522R0R0 | A0A522R0Q8 | A0A522R0Q3 | A0A522R0P9 |
| ***Sinorhizobium mexicanum* strain ITTG R7** | 375549 | A0A859QIV6 | A0A859R0Y0 | A0A859QZ22 | A0A859R570 | A0A859QVQ8 | A0A859QW65 | A0A859QRE9 | A0A859QMWZ |
| ***Hyphomicrobiales* bacterium strain SCN18_10_11_15_R4_B_69_13** | 1909294 | A0A8I1P1S7  A0A8I1P530 | A0A8I1P690 | A0A8I1P1S0 | A0A8I1TIF2 | A0A8I1THA1 | A0A8I1TKX7 | A0A8I1TFV6 | A0A8I1TET4 |
| ***Chelativorans petroleitrophicus* strain SCAU2102** | 2975484 | A0A9X2XAV7 | A0A9X3B0R8 | A0A9X2XAH4 | A0A9X2XB67 | A0A9X3B7M2 | A0A9X3BAF7 | A0A9X3V7T3 | A0A9X3B0I0 |
| ***Paracoccus onubensis* strain L29414** | 1675788 | A0A418T7P6 | A0A418T7C3 | A0A418T7C2 | A0A418T7H0 | A0A418T7E5 | A0A418T7D2 | A0A418T7P8 | A0A418T7C4 |
| ***Paracoccus methylarcula* strain H1** | 72022 | – | – | A0A3R7NEB5 | – | A0A422R1M1 | – | – | – |

**Table S6.** Uniprot sequence accession codes of ComC, SlcC and SuyAB homologues of *P. wurundjeri* strain Merri biomineralization proteins.

|  | **Taxon ID** | **ComC** | **SlcC** | **SuyA** | **SuyB** |
| --- | --- | --- | --- | --- | --- |
| ***Paracoccus alkanivorans* strain 4-2** | 2116655 | A0A3M0MF96 | A0A3M0MKS2 | A0A3M0MZ42 | A0A3M0MES7 |
| ***Rhizobiales* bacterium strain 65-79** | 1895815 | – | – | – | – |
| ***Rhizobiaceae* bacterium strain FW021_bin.52** | 1913961 | A0A522R0R3 | A0A522R0P3 | – | A0A522R0P0 |
| ***Sinorhizobium mexicanum* strain ITTG R7** | 375549 | A0A859QZ08 | A0A859QS75 | A0A859QGT8 | A0A859R0Z4 |
| ***Hyphomicrobiales* bacterium strain SCN18_10_11-15_R4_B_69_13** | 1909294 | A0A8I1P3J4 | – | – | – |
| ***Chelativorans petroleitrophicus* strain SCAU2101** | 2975484 | A0A9X2XAT1 | A0A9X2X9Q3 | A0A9X2XBV0 | A0A9X2XCG5 |
| ***Paracoccus onubensis* strain L29414** | 1675788 | A0A418T7A4 | A0A418T7D0 | A0A418T7A7 | A0A418T7M7 |

**References**

(1) Denger, K.; Cook, A. M. Racemase activity effected by two dehydrogenases in sulfolactate degradation by *Chromohalobacter salexigens*: purification of (*S*)-sulfolactate dehydrogenase. *Microbiology* **2010**, *156*, 967-974.

(2) Denger, K.; Weiss, M.; Felux, A. K.; Schneider, A.; Mayer, C.; Spiteller, D.; Huhn, T.; Cook, A. M.; Schleheck, D. Sulphoglycolysis in *Escherichia coli* K-12 closes a gap in the biogeochemical sulphur cycle. *Nature* **2014**, *507*, 114-117.

(3) Felux, A. K.; Spiteller, D.; Klebensberger, J.; Schleheck, D. Entner-Doudoroff pathway for sulfoquinovose degradation in *Pseudomonas putida* SQ1. *Proc. Natl. Acad. Sci. USA* **2015**, *112*, E4298-4305.
